# Multiple brain activation patterns underlying successful visual short-term memory across the adult lifespan

**DOI:** 10.1101/2024.10.08.617160

**Authors:** Michelle G. Jansen, Alireza Salami, Fernando Ruiz Martínez, Daniel J. Mitchell, Joukje M. Oosterman, Linda Geerligs

## Abstract

Cognitive task performance can be supported through multiple brain activation patterns, a concept referred to as brain degeneracy. We used a novel approach to consider brain degeneracy during a visual short-term memory task (VSTM) across the adult lifespan in the cross-sectional Cam-CAN study (n = 113, 23-87 years old). Our main goal was to identify subgroups of participants whose VSTM performance was characterized by distinct brain activation patterns. First, we identified seven brain modules that responded similarly to the VSTM task across participants, and resembled previously identified functional networks. Subsequently, latent profile analysis revealed four distinct subgroups of participants. Each subgroup was characterized by different recruitment patterns of these brain modules, predominantly in the frontal control module (FCM), visual module (VM), and default mode module (DMM). Subgroups did not differ in demographics or task performance. However, associations between brain activity and performance varied across subgroups, particularly in the FCM, suggesting that individuals may use different cognitive operations to perform the VSTM task. Further analyses revealed group differences in white matter integrity, mostly in the uncinate fasciculus, suggesting that individual differences in structural brain properties may shape the different brain activation patterns. Altogether, our study contributes to our understanding of how multiple brain activation patterns could underlie cognitive performance.

## 1 Introduction

Over the past decades, research sought to elucidate how the brain’s functional and structural architecture gives rise to cognitive functioning (Finn et al., 2023; Genon et al., 2022; Park & Friston, 2013; Poldrack, 2006). Nevertheless, this relationship is far from straightforward. Not only do brain measures typically account for only a small portion of the variance in cognitive functioning (Hedden et al., 2016), but there are also several contradictory findings in terms of which neural mechanisms are adaptive or maladaptive for cognitive performance. For example, age-related increases in prefrontal activation have been linked to both superior and inferior performance in older adults (Grady, 2012; Nyberg et al., 2020). The complex nature of brain-cognition associations is further emphasized by previous studies showing that the extent to which brain structure or function relates to cognitive performance varies substantially across individuals (Lindenberger, 2014; Lövdén et al., 2018; Patel et al., 2022; Salami et al., 2018).

To gain a better understanding of how cognitive functions are supported throughout the lifespan, we may need to revisit the common assumption that most individuals exhibit similar patterns of neural activity to perform a certain cognitive task (one-to-one mapping) (Seghier & Price, 2018; Westlin et al., 2023). Conversely, brain degeneracy suggests that a specific task could be executed through multiple neural systems (many-to-one mapping) (Edelman & Gally, 2001; Noppeney et al., 2004; Price & Friston, 2002). Degeneracy is evident in many neural systems, including those supporting cognitive functions (Kelso, 2012; Noppeney et al., 2004). In line with this, previous studies have identified subgroups of individuals showing unique neural activity patterns while performing the same task (Cerliani et al., 2017; Kherif et al., 2009; Seghier et al., 2008). Interestingly, this neural variability was associated with the use of different strategies to complete the task (Kherif et al., 2009; Noppeney et al., 2006), despite similar levels of task performance (Fischer-Baum et al., 2018; Noppeney et al., 2006). This illustrates how multiple neural routes can lead to successful cognitive performance. However, if we focus on similarities across individuals these possibilities are overlooked. Furthermore, analyses aimed at capturing individual differences may miss meaningful associations when structured sources of variability exist. For example, if two subgroups within the same sample show opposing brain-behaviour associations, these effects may cancel out at the group level (Westlin et al., 2023). Recognizing and considering this heterogeneity is therefore essential to understand brain degeneracy and what it might mean for how we study the associations between brain and cognition.

Building on the concept of many-to-one mapping, it has been suggested that secondary activation repertoires may be prompted when the primary circuit is unable to sustain successful task completion (Mason et al., 2015; Noppeney et al., 2004; Price & Friston, 2002). For instance, studies using transcranial magnetic stimulation showed that younger adults adapt to temporary functional lesions by recruiting alternative neural circuits (Lee et al., 2003; Pascual-Leone et al., 2005). Furthermore, differential recruitment of brain regions during task performance has been related to functional recovery after acquired brain injury (Nudo, 2013). In the context of normal ageing, alterations in brain activity have been linked to the degradation of major white matter tracts (Burzynska et al., 2013; Lucas et al., 2018), and localized grey matter loss (Kalpouzos et al., 2012; Salami et al., 2012; Salami et al., 2014). Although this involves functional deterioration, older adults are able to compensate for impaired brain function by recruiting additional brain regions, particularly in the frontal lobes (Cabeza et al., 2002; Grady, 2012; Park & Reuter-Lorenz, 2009). In contrast, structural preservation in older individuals (i.e., brain maintenance; Nyberg et al., 2012) has been related to neural activation patterns more closely resembling those typically observed in younger adults (Cabeza et al., 2018; Nyberg et al., 2012). Together, these findings stress that it is important to consider interindividual variability in brain structure to understand why different neural systems may be recruited during task performance.

In the present study, we aimed to investigate brain degeneracy across the adult lifespan in the context of visual short-term memory (VSTM), using cross-sectional data from the Cam-CAN project (*n* = 122, 23-87 years old). VSTM is particularly interesting in the context of degeneracy, given that high performance can be achieved via different strategies (Ritakallio et al., 2024). VSTM refers to our ability to maintain visual information and is crucial for many other cognitive functions (Luck & Vogel, 2013; Pearson & Keogh, 2019). The term VSTM emphasizes maintenance, without additional manipulation of information, which is typically referred to as visual working memory. The precise neural mechanisms underlying successful VSTM performance are not fully understood, partially due to inconclusive findings in prior research that did not consider the possibility of different brain activation patterns (Pearson and Keogh, 2019). Our primary aim was therefore to identify subgroups of participants whose VSTM performance was characterized by distinct brain activation patterns. We first identified modules of brain regions that showed similar responses across participants during the VSTM task. Next, we applied latent profile analysis to identify sub-groups of participants that demonstrated differential recruitment patterns of the previously identified brain modules. To better understand the variability in brain activity during the VSTM task, we characterized the most significant differences in brain activity across subgroups.

Furthermore, we sought to elucidate factors contributing to the observed subgroup differences in brain activity patterns. Specifically, we investigated the extent to which these subgroups differed in demographic variables (i.e., age, sex, educational attainment) and task performance. We were particularly interested in the role of age, as this factor was previously related to both performance and neural correlates of the VSTM task (Lugtmeijer et al., 2023; Mitchell & Cusack, 2018; Morcom & Henson, 2018; Sander et al., 2012). For example, if age-related differences in brain function were the largest source of inter-individual variability in brain activation patterns during the VSTM task, we expected to identify subgroups that differed in their age and task performance. Conversely, if age-invariant inter-individual differences were most prominent, we expected to see participant subgroups that were roughly age-matched. Additionally, we investigated whether structural brain characteristics, such as grey matter volumes and white matter integrity, would explain variations in brain activity among the subgroups, as reflected by subgroup differences in these measures.

Lastly, we assessed associations between brain activation profiles and task performance across all participants, and separately within each subgroup. For example, while no association might be observed in the overall sample, increased activity in brain module X might have been beneficial for performance for subgroup Y but detrimental for subgroup Z. Such unique associations would not only provide further evidence for the presence of different neural routes to facilitate successful VSTM performance, but also potentially suggest cognitive strategies that were used to succeed.

## 2 Methods

### 2.1 Participants

This study included data from 122 participants (59 female, 48.4%) who were aged 23–87 (mean age 53.9, SD = 18.1) from the healthy, population-derived cohort tested in Stage III of the Cam-CAN project (Shafto et al., 2014; Taylor et al., 2017).

A complete overview of the study procedures and inclusion criteria have been described previously (Shafto et al., 2014; Taylor et al., 2017). Briefly, in Stage I, participants engaged in home-based interviews where demographic information, cognitive measures, and other measures of health (e.g., mental and physical health) were collected. In Stage II, participants engaged in three testing sessions, including cognitive and behavioural assessments, structural MRI (e.g., T1-weighted imaging and diffusion-weighted imaging), functional MRI (e.g., resting-state), and MEG recordings. During Stage III, among other things, structural MRI (e.g., T1-weighted images) and task-related functional MRI (fMRI) data during a VSTM task were acquired. In this manuscript, we use the demographic information from Stage I, the diffusion-weighted images from Stage II, and the T1-weighted images and VSTM fMRI paradigm from Stage III. Participants were native English-speakers with normal or corrected-to-normal vision and hearing and had no neurological disorders. Additional exclusion criteria were history of drug or alcohol abuse, and poor hearing or vision.

The study was conducted in accordance with ethical approval obtained from the Cambridgeshire 2 (now East of England - Cambridge Central) Research Ethics Committee. All participants gave written informed consent prior to participation.

We excluded a total of 9 participants from our analyses due to poor VSTM task performance (*n* = 8) or data quality issues (*n =* 1), leaving a final sample of 113 participants (see below for more details).

### 2.2 Data acquisition

#### 2.2.1 Demographic information

Demographic information, including age, sex, and educational attainment, was acquired during home-based interviews during Stage I of the Cam-CAN project. Educational attainment in this study was based on the highest level of completed education (i.e., categorized in 1 = no education above the age of 16 years old; 2 = GCSEs or equivalent; 3 = A levels or equivalent; 4 = university levels or equivalent).

#### 2.2.2 MRI data

MRI data was acquired on a 3T Siemens TIM Trio System scanner at the MRC Cognition Brain and Sciences Unit, Cambridge, UK. During each of three VSTM task runs, functional data were acquired using a multi-echo, T2*-weighted EPI sequence (TR = 2000 milliseconds, TE = 12 milliseconds, 25 milliseconds, 38 milliseconds, flip angle = 78 degrees, 32 axial slices of thickness of 3.7 mm with an interslice gap of 20%, FOV = 294mm × 294 mm, voxel-size = 3 mm × 3 mm × 3.48 mm). Each volume had 34 axial slices (acquired in descending order), slice thickness of 2.9 mm, with an interslice gap of 20%. The multiple echoes were combined by computing their average weighted by their estimated T2* contrast. The number of scans acquired varied per run, depending on reaction time, between 294 and 349.

A high-resolution 3D T1-weighted structural image was acquired using a Magnetization Prepared Rapid Gradient Echo (MPRAGE) sequence, with the following parameters: TR = 2250 ms; TE = 2.99 ms; TI = 900 ms; flip angle = 9 degrees; FOV = 256 × 240 × 192 mm; voxel size = 1 mm isotropic; GRAPPA acceleration factor = 2. This T1 image was missing for two of the participants in this sample, however for these participants T1 images were acquired during the scanning session that took place during Stage II of Cam-CAN, approximately 1-3 years earlier.

Diffusion-weighted images (DWIs) were acquired with a twice-refocused spin-echo sequence during Stage II of Cam-CAN. Thirty diffusion gradient directions were obtained for each of two b-values: 1000 and 2000 s/mm2, plus three images acquired with a b-value of 0. These parameters were optimized for estimation of the diffusion kurtosis tensor and associated scalar metrics (Shafto et al., 2014). Other DWI parameters were TR = 9100, TE = 104 ms, FOV = 192 mm × 192 mm, voxel size 2 mm isotropic, 66 axial slices, with acquisition time 10 minutes and 2 seconds.

#### 2.2.3 Visual short-term memory (VSTM) task

During fMRI scanning, participants performed a VSTM paradigm (see **Figure 1)**, based on Emrich et al. (2013). A detailed description of the task is provided in Lugtmeijer et al. (2023).

**Figure 1:**
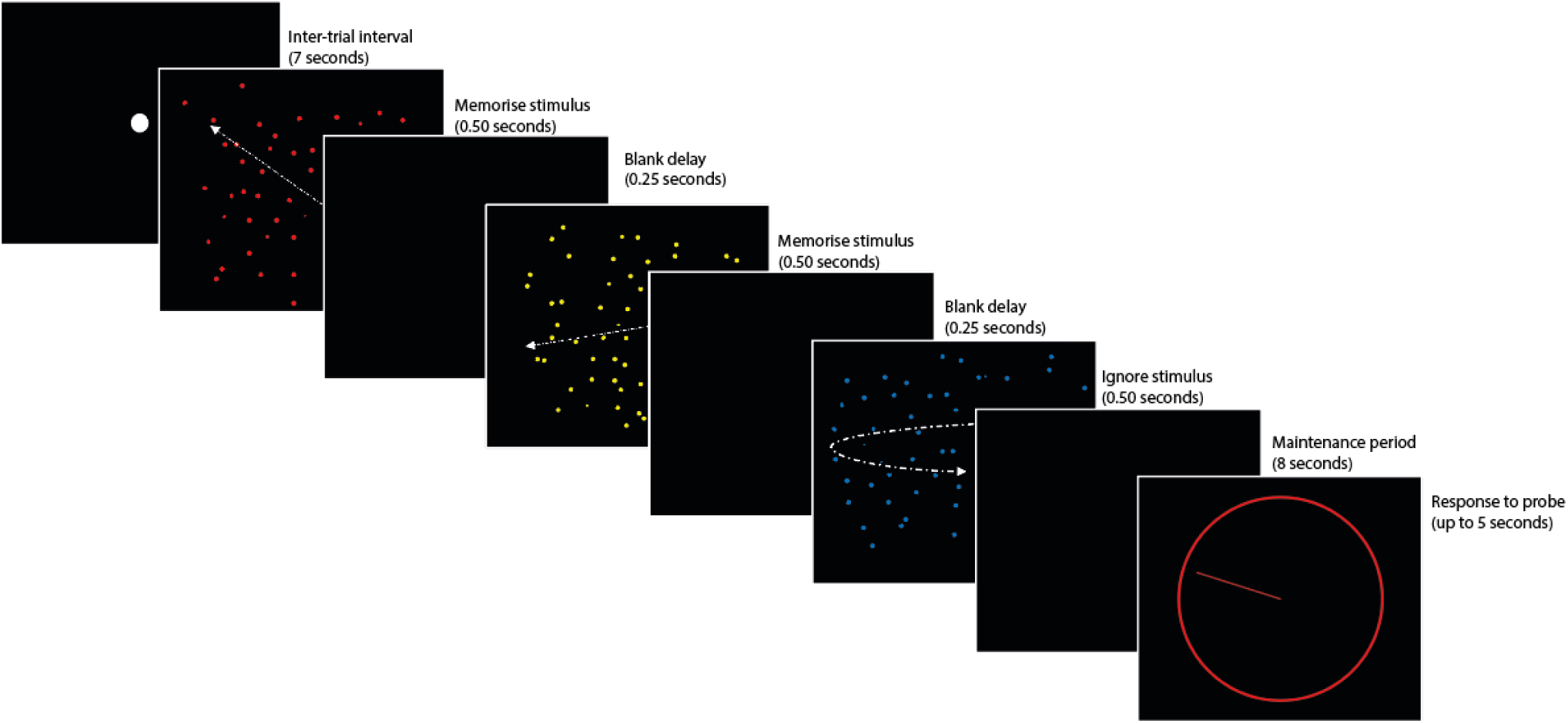
Adapted from Morcom and Henson (2018). An example of the VSTM task. Trials began with a fixation dot for 7 seconds. On each trial, three moving dot displays were then presented in red, yellow, and blue for 500 ms each (250 ms in between). After the last display, the screen was blank for an 8 second maintenance period and then a probe display was presented. A coloured circle indicated which dot display participants should recall (red, yellow, or blue). Participants had up to 5 seconds to adjust a pointer until its angle matched the motion direction of the probed display.

Briefly, three displays containing different coloured moving dots (red, yellow and blue) were presented sequentially on each trial. Each trial began with a grey fixation dot for 5 seconds. Subsequently, the fixation dot brightened for 2 seconds and was followed by the coloured dot displays. Each coloured dot display was shown for 500ms, with a 250ms black screen in between. As a manipulation of the number of items in memory, trials could contain one, two or three arrays of dots moving in a single direction (load 1, 2, and 3, respectively).

Participants were instructed to remember the direction of motion for dot displays in which all dots moved in a single direction, and to ignore any dot displays that rotated around the fixation location (the encoding phase). As three displays were shown in each trial, the memory load was manipulated by varying the number of rotating dots in relation to the number of moving dot displays. After the third coloured dot display, there was a blank display of 8 seconds (the maintenance phase), followed by a probe display. The probe display consisted of a coloured circle that indicated which of the three dot displays (red, yellow, or blue) the participant should recall. Participants had up to 5 seconds to adjust a pointer on the circle until it matched the direction of motion of the probed dot display (the response phase).

### 2.3 VSTM performance

Several factors can limit VSTM performance. For example, memory imprecision (e.g., uncertainty between 80 or 85 degrees), random guessing (e.g., when forgetting an item entirely), and reporting an incorrect item (mis-binding, e.g., reporting the angle of the red dots instead of the blue dots). To distinguish these different sources of error, we applied a three-component mixture model (Bays et al., 2009). A complete description of the application of the mixture model to the VSTM task is provided in previous work (Mitchell et al., 2018; Lugtmeijer et al., 2023).

For our subsequent analyses, we used the estimates of a memory item’s precision (the inverse of the standard deviation in degrees) and the number of items (directions) that were stored in memory (K). Eight participants in our sample showed very poor performance for memory of a single item (>30 degrees deviation on average). This suggests that they may not have understood the task instructions and were therefore excluded from further analyses. Since there was a ceiling effect for the number of items in memory at load 1 (mean = 0.96, SD = 0.06) and to some extent at load 2 (mean = 1.76, SD = 0.24), we decided to focus on performance at load 3 (mean = 2.40, SD = 0.46). The distribution of scores for the number of items in memory at each load is shown in **Supplementary Figure 1**.

### 2.4 MRI pre-processing and first level analyses

All MRI data were pre-processed using SPM 12 (http://www.fil.ion.ucl.ac.uk/spm) and “automatic analysis” software (Cusack et al., 2015). The T1 and T2 weighted structural images were rigid body registered with a MNI template, bias corrected and segmented into six tissue classes using the newer segment protocol in SPM 12 (Ashburner & Friston, 2005). Non-linear transformation using DARTEL normalization was performed to match a grey matter template created from the Cam-CAN stage III sample. The functional images were spatially realigned and interpolated to correct for differences in acquisition times, and co-registered to the structural image using rigid body transformations. Both grey matter volume images and functional images were transformed to MNI space using the warps and affine transformation estimated from the structural images. For grey matter volume images, this was done while preserving the total amount of signal from each region in the images. The functional images were resliced to 3×3×3 mm voxels, while grey matter volume images were resliced to 1.5×1.5×1.5 mm voxels and both were smoothed with an 8 mm full-width half-maximum (FWHM) Gaussian kernel. For the two participants with missing T1-weighted scans, normalization and co-registration were performed based on the T1 images collected during stage II of the Cam-CAN project. One participant was excluded from further analyses due to data quality issues.

For each participant and voxel, we fit a general linear model (GLM) with three phases of each trial per run: encoding, maintenance, and response. The encoding phase was modelled as an epoch of 2 seconds starting from the beginning of the first dot display movement. The maintenance phase was modelled as an epoch of 8 seconds (i.e., the duration of the blank display that was shown after all probes were displayed). The response phase was modelled as an epoch of the time between the probe onset and response. These three components were each split into three regressors: one for each load. Six additional regressors were added per run, representing the motion parameters estimated in the realignment stage. The means of each run were also modelled separately. The main contrast of interest in this study was the difference between the highest and the lowest memory load during the maintenance period (load 3 vs. load 1). This contrast allows us to equate the amount of visual information that was presented just before the maintenance period as well as the motor response that was required just after, while maximizing the difference in memory load.

### 2.5 Identifying modules with similar inter-individual variability

We first set out to identify modules with similar inter-individual variability across a smaller subset of brain modules, which involved several dimensionality reduction steps. This was done to make the analyses in later stages more computationally efficient and interpretable, particularly for the latent profile analyses (LPA; see *2.7 Structural brain characteristics* for details). LPA are typically performed on a limited number of input variables to avoid convergence issues and unstable solutions (Spurk et al., 2020).

First, we grouped voxels into a set of small regions of interest (ROI). Specifically, we used the 748 (out of 840) regions from the Craddock atlas (2012) that had sufficient coverage in our recent paper (Geerligs et al., 2015), allowing us to use the network labels defined in our previous study. To select ROIs that might be involved in task performance in a subset of participants, we first extracted the average t-values for the contrast of interest (i.e., load 3 vs. load 1 during the maintenance period) across all voxels in each ROI for each participant. To further reduce the dimensionality of the data, we only included ROIs that showed increased or reduced activation in load 3 versus load 1 in at least 10% of the participants at a liberal significance threshold (T > 1.96, p < 0.05). It should be noted that this threshold was not empirically derived; we chose this threshold for a balance between excluding regions likely driven by noise and including those that may be relevant for task performance in a meaningful subset of individuals (i.e., 10% of our sample corresponding to N±12). This procedure resulted in the final inclusion of 418 ROIs.

For each included ROI, we extracted the average beta-values. Subsequently, we calculated the correlation values between ROIs for load 3 vs. load 1 across participants (i.e., ROI-by-ROI correlation matrix). A high correlation coefficient between two ROIs suggests that these regions show similar inter-individual variability in load-dependent modulation of the BOLD response, and therefore may contribute to the same brain module.

In the next step, we used the Brain Connectivity Toolbox to identify groups of ROIs, or brain modules, that showed similar brain activity patterns across participants (Rubinov & Sporns, 2010). Specifically, we applied consensus partitioning based on the Louvain modularity algorithm (Bassett et al., 2013; Lancichinetti & Fortunato, 2012). We opted for this algorithm as it is one of the most widely used methods in network neuroscience, largely due to its computational efficiency and its ability to identify modules with high modularity (i.e., high within-module and low between-module connectivity) (Blondel et al., 2008; Rubinov & Sporns, 2010). The ROI-by-ROI correlation matrix was used to create an initial partition of the ROIs into distinct modules with the Louvain modularization algorithm (Blondel et al., 2008), and was further refined using an optimization procedure (Sun et al., 2009). This partitioning procedure was repeated 500 times, and all the repetitions were assembled into an ROI-by-ROI agreement matrix, where each number indicates the proportion of repetitions in which a pair of ROIs were assigned to the same module. The ROI-by-ROI agreement matrix was then used as an input for consensus partitioning, where a permuted version of the agreement matrix was used for thresholding. This was repeated until the algorithm converged into a matrix of ones and zeros, indicating that a single partition was reached.

The complete partitioning procedure was performed for multiple values of the resolution parameter γ, ranging from 1 to 1.5 with steps of 0.05. The γ parameter is a penalty term that biases the size of the communities to be found. Smaller values of γ will result in larger modules, while higher values will result in smaller modules. An γ value of 1 resulted in 3 large modules, providing a very coarse parcellation and limited information about participants’ activation profiles. Conversely, an γ value of 1.5 already resulted in 49 modules. Therefore, we did not explore a wider range of this parameter space.

Upon inspection of the resulting modules, we noted that the identified brain modules appeared highly similar to traditional functional networks. We therefore selected the partition with the highest resemblance to a previously defined set of age-representative networks (Geerligs et al., 2015b), as measured with the adjusted mutual information (aMI), as our final set of brain modules. This allowed us to select a modular structure that aligned with known network organization across the adult lifespan, improving interpretability and comparability with previous work. The optimum we identified was at γ = 1.25, resulting in 13 distinct modules (aMI = 0.45). However, as many modules consisted of only one ROI or a few ROIs, we only included modules with at least 15 ROIs in our analyses. This resulted in the exclusion of 6 brain modules, comprising 28 ROIs in total. The final set of 7 brain modules was used for further analyses (see “Results”).

Within each brain module and for each participant, we computed the average brain activity for load 3 vs. load 1. However, we observed strong correlations between single-module activity and the average brain activity values across all brain modules (ranging from r=0.39 to 0.77). This is consistent with prior research which demonstrated that overall network responsivity, rather than module-specific activation patterns, were related to age and task performance (Samu et al., 2017). To be able to look beyond this general factor in our subsequent analyses, we computed the residual activity within each brain module after regressing out the mean activity across the 418 ROIs (i.e., averaged across all participants). The proportion of variance explained in the activity of each ROI by the mean activity across ROIs ranged from 0.61 to 0.75, as reflected by the R² values from these regression models.

However, because residuals are centered around zero, they no longer reflect the overall magnitude of activity in each brain module (i.e., whether a module was generally more or less engaged during the task). To be able to retain information about the engagement of each network during the task, the mean activity across all subjects was added back after computing the residuals. Importantly, this adjustment does not affect the statistical comparisons in our subsequent analyses; rather, this restores interpretability by allowing the values to reflect relative differences in brain activity.

### 2.6 Brain activity profiles

To identify latent subgroups of participants characterized by different patterns of brain activity, we applied LPA using the Mclust package in R (Scrucca et al., 2016). Briefly, LPA is a data-driven Gaussian mixture modelling method to identify hidden subgroups in a population based on multiple indicator variables (Vermunt & Magidson, 2002), such as brain activity across several modules. A Gaussian mixture model assumes a multivariate Gaussian distribution for each subgroup. Therefore, every observation (here, participant) will have a specific probability of belonging to a specific subgroup, as specified by a probability density function determined by the average, the variance, and the covariance of each of the subgroup distributions. We opted for LPA because it is able to model different data shapes, including elliptical, skewed, or dispersed shapes, making it flexible for real-world, heterogeneous data (Lövdén et al., 2018; Salami et al., 2019; Spurk et al., 2020).

We applied LPA using eight indicators: the residual activity in each of the seven brain modules and one variable capturing the overall responsivity across all networks (i.e., 122 rows per subject x 8 columns). This allowed us to look at module-specific variations in responsivity in addition to the overall responsivity across brain modules. We restricted the model space to models that do not allow for covariance between modules within subgroups of participants (i.e., models with a spherical or diagonal distribution of the within-group covariance matrix). This way, our model assumes that any correlations between modules must be caused by participants with different brain activity profiles (similar to Lövdén et al., 2018).

To select the optimal model and class solution (i.e., number of subgroups), we used the Bayesian Information Criterion (BIC), Integrated Completed Likelihood (ICL) and bootstrapped Likelihood Ratio Test (LRT). In this context, larger BIC and ICL values indicate a better model fit (Scrucca et al., 2016).

### 2.7 Structural brain characteristics

To investigate subgroup differences in grey matter, we extracted grey matter volumes for each of the 418 ROIs that were included in our main analysis and subsequently averaged the grey matter volumes within each module. The two participants with the missing T1-scan at Stage III were excluded from this analysis, resulting in a sample of 111 participants for the grey matter data.

To investigate subgroup differences in white matter microstructure, we used the diffusion data to estimate mean kurtosis. Mean kurtosis captures the degree to which water diffusion deviates from a Gaussian distribution within a voxel, and is influenced by changes in organelles, cell membranes and the ratio of intracellular to extracellular water compartments (Falangola et al., 2008). It provides a sensitive metric to determine how aging affects the complexity of white matter microstructure, and typically decreases in late-life (Benitez et al., 2018).

Diffusion data were pre-processed and analysed as described in a previous paper (Geerligs et al., 2018). Briefly, the data were co-registered, normalized to MNI space and smoothed with a 1-mm FWHM Gaussian kernel to reduce residual interpolation errors. Subsequently, we calculated the average mean kurtosis values for 18 tract-based regions of interest defined by the Johns Hopkins University white matter tractography atlas (Hua et al., 2008). These included the uncinate fasciculus, superior longitudinal fasciculus, inferior fronto-occipital fasciculus, anterior thalamic radiations, forceps minor, forceps major, cerebrospinal tract, the inferior longitudinal fasciculus, ventral cingulate gyrus, and the dorsal cingulate gyrus. All tracts were split into their left and right hemisphere sections, except for the forceps minor and forceps major.

Participants with severe head motion during the DWI scan, as revealed by an automated striping detection method on the diffusion data, were excluded (Neto Henriques and Correia, 2015). In addition, for each tract, participants with mean kurtosis values of 3 SDs above or below the group average were excluded. This resulted in a final sample of 96 participants for all tracts, except for the inferior longitudinal fasciculus (n = 95), superior longitudinal fasciculus (n = 95), cerebrospinal tract (n = 94), and ventral cingulate gyrus (n = 86).

### 2.8 Statistical tests

All statistical tests were performed using R (version 4.3.1).

Given that overall responsivity (i.e., average activity across all included ROIs) explained most of the variability in brain module activity, we first explored how this factor related to age and task performance using Pearson correlations. To investigate whether associations between overall responsivity and task performance were age-independent, we additionally calculated the partial correlations between these two variables while accounting for age.

Next, we performed several between-group comparisons to better understand the diverse brain activity profiles associates with the VSTM task that were identified with the latent profile analysis. To assess which brain modules showed the most pronounced differences across subgroups we statistically compared brain module activity between groups. We also tested whether subgroups differed in demographic characteristics (age, sex, educational attainment), VSTM task performance (the number and precision of items in memory at load 3), grey matter volumes (i.e., average grey matter volumes in each of the brain modules), and white matter microstructure (i.e., average mean kurtosis in each of the 18 tracts). These comparisons were performed to understand which factors contributed to the differences in brain activity profiles.

We applied a two-step approach to determine between-group differences. First, Welch’s ANOVA was used to determine if there were differences between subgroups in the variable of interest. This was used instead of the regular Fisher’s one-way ANOVA because of unequal sizes and variances between groups. Second, each significant ANOVA was followed by pairwise Welch’s t-tests between groups (i.e., using non-pooled standard deviations to consider the unequal variances between groups). Chi-squared tests were used in case of categorical data (i.e., comparing sex or educational attainment across groups). As educational attainment had fewer than five observations in certain categories, this variable was split into a dichotomous category prior to performing the comparisons (university level vs. other educational categories), however for descriptive purposes all categories are reported.

Analyses on grey matter volumes were corrected for intracranial volume. All analyses were repeated while additionally correcting for the effects of age, except for the comparisons on demographic characteristics. Specifically, we regressed the covariate(s) on the variable of interest and used the residuals of that regression for the ANOVA and subsequent t-tests.

To further investigate the extent to which different neural routes facilitated successful VSTM performance, we investigated the associations between brain module activity and task performance across all participants, and separately within each subgroup. If certain associations are present in one subgroup but not in another, this could suggest that these subgroups relied on different cognitive operations to successfully complete the task. Similarly, if an association is present within a subgroup but not at the group level, this illustrates how subgroup-specific effects may be obscured in whole-sample analyses. To test this, we calculated Pearson correlations between brain module activity and task performance in each of these groups, adjusting for the effects of age (i.e., partial correlations). Difference between coefficients were additionally statistically compared using Fisher’s z-tests.

We used false discovery rate (FDR) corrections (Benjamini & Hochberg, 1995) to adjust for multiple comparisons across different brain modules, white matter tracts or pairwise comparisons between-groups, where appropriate. In addition to reporting (FDR-corrected) p-values, we computed Bayes Factors (BF) for the alternative hypothesis using the *BayesFactor* package (https://cran.r-project.org/web/packages/BayesFactor/index.html), with default settings. Given the exploratory nature of our analyses, effects were considered as strongly supported in case of both FDR-adjusted p < 0.05 and BF > 3 (Buades-Rotger et al., 2023; Driessen et al., 2018), while those only surpassing uncorrected p < 0.05 are interpreted with caution. In addition, we calculated Bayesian R^2^ to evaluate the fit of each model (Gelman et al., 2019), using the *performance* package (https://easystats.github.io/performance/index.html).

## 3 Results

### 3.1 Brain modules with similar brain activity patterns across participants

We identified seven distinct modules of brain regions with similar patterns of inter-individual variability in terms of their brain activity while performing the VSTM task (**Figure 2**). The brain activity metric used to define these modules (and used in all subsequent analyses) was based on the contrast of load 3 vs. load 1 during the maintenance period.

**Figure 2:**
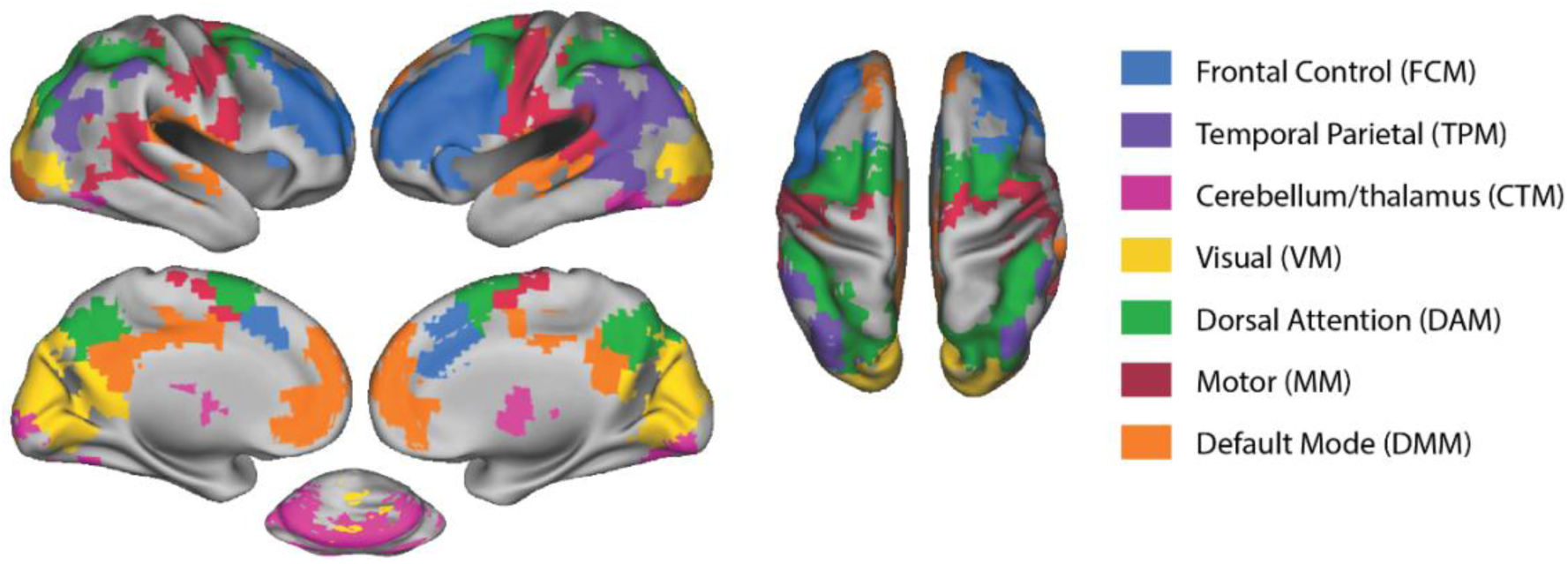
An illustration of the seven brain modules identified. For visualization purposes, data were projected to surface space using the Caret toolbox (Van Essen et al., 2001).

The modules included a frontal control module (FCM), which overlapped largely with the traditional fronto-parietal control network. This module encompasses the lateral prefrontal cortex and the anterior cingulate but did not include parietal areas. A distinct temporo-parietal module (TPM) included parietal areas that typically fall within the default mode and fronto-parietal control network, predominantly in the left hemisphere, but also included temporal areas. The cerebellum and thalamus module (CTM) covered the cerebellum and thalamus, and the visual module (VM) included the occipital cortex. Another module overlapped with the dorsal attention network, covering the frontal eye fields and the intraparietal sulcus, which we refer to as the dorsal attention module (DAM). The motor module (MM) covered the bilateral precentral gyrus but also extended into the right posterior temporal cortex. Lastly, one module largely overlapped with the default mode network, covering the medial prefrontal cortex, precuneus, and temporal areas, which we refer to as the default mode module (DMM).

### 3.2 Overall responsivity increases with age and accounts for module-specific brain activity

As previously described (Methods 2.5), the average activity across all included ROIs explained a substantial proportion of the variance in brain activity within each module (R^2^ = 0.61-0.75).

We explored how the overall responsiveness related to age and task performance. Average activity across all brain modules increased with age (r = 0.37, 95% CI = [0.19; 0.52], uncorrected p < 0.001, BF = 338.96). However, we did not observe significant associations between overall activity and memory precision (r = 0.04, uncorrected p = 0.688, BF = 0.15) or number of items in memory (r = −0.07, uncorrected p = 0.460, BF = 0.18).

### 3.3 Latent profile analysis identifies subgroups with distinct activation signature across modules

To look beyond overall responsivity, this factor was regressed out of the seven identified brain networks. To account for all variance in brain activity, the LPA grouped participants based on the residual brain activity within each brain network, in addition to the overall responsivity across all ROIs. The distributions of BIC and ICL values for the different latent profile models across a wide range of classes are shown in **Supplementary Figure 2 and 3**. Both the BIC and ICL suggested that the model with equal shape, variable volume, and assumed diagonal covariance (“VEI” model) provided the most optimal fit. Subsequently, to select the number of classes, we compared the BIC and ICL values, as well as the bootstrapped LRT results. The BIC and ICL both indicated that the solution with 4 classes provided the most optimal fit (BIC = −1028, ICL = - 1047.89). Further LRT analyses resulted in a non-significant result only at 8 classes (p = 0.422), suggesting that a model with 7 classes had the best model fit (**Supplementary Table 1**). However, the BIC and ICL continued to show the most optimal fit at 4 classes. As we aimed for a parsimonious solution, we therefore opted for the solution with 4 subgroups (n = 35, n = 24, n = 42, and n = 12 in groups 1-4, respectively).

An overview of the brain module activation profiles across subgroups is displayed in **Figure 3**. To investigate which networks were the strongest drivers of the subgroup differences, we statistically compared the brain activity profiles across subgroups **(Table 1).** It should be noted that the LPA was designed to produce subgroups with distinct brain activity profiles. Therefore, these significance values are only meaningful in relation to each other as a comparison of the relative contributions of different modules and subgroups, but not in isolation. A complete overview of the between-group statistics for the brain module activity is depicted in **Supplementary Table 2**.

**Figure 3:**
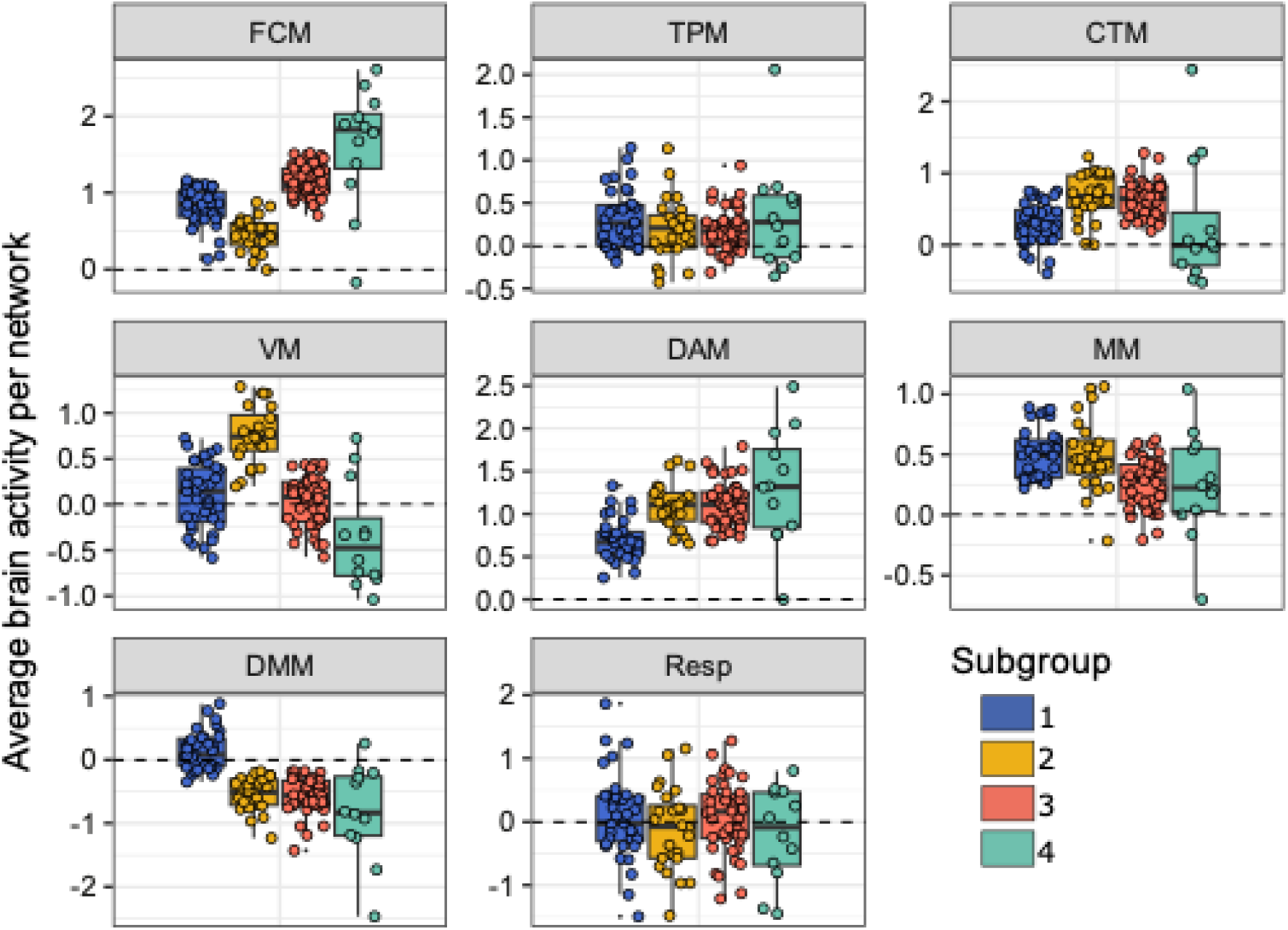
For each of the four identified subgroups this figure shows the mean-regressed average brain activity in each of the seven identified brain networks as well as the overall responsivity across all ROIs (Resp). The most influential brain modules were the FCM, VM, DAM, and DMM, as they showed strong evidence for subgroup differences (BF > 100, q < 0.001) and substantial explained variance (Bayesian R² ≥ 0.26). FCM = frontal control module; TPM = temporal parietal module; CTM = cerebellum/thalamus module; VM = visual module; DAM = dorsal attention module; MM = motor module; DMM = default mode module.

**Table 1.**
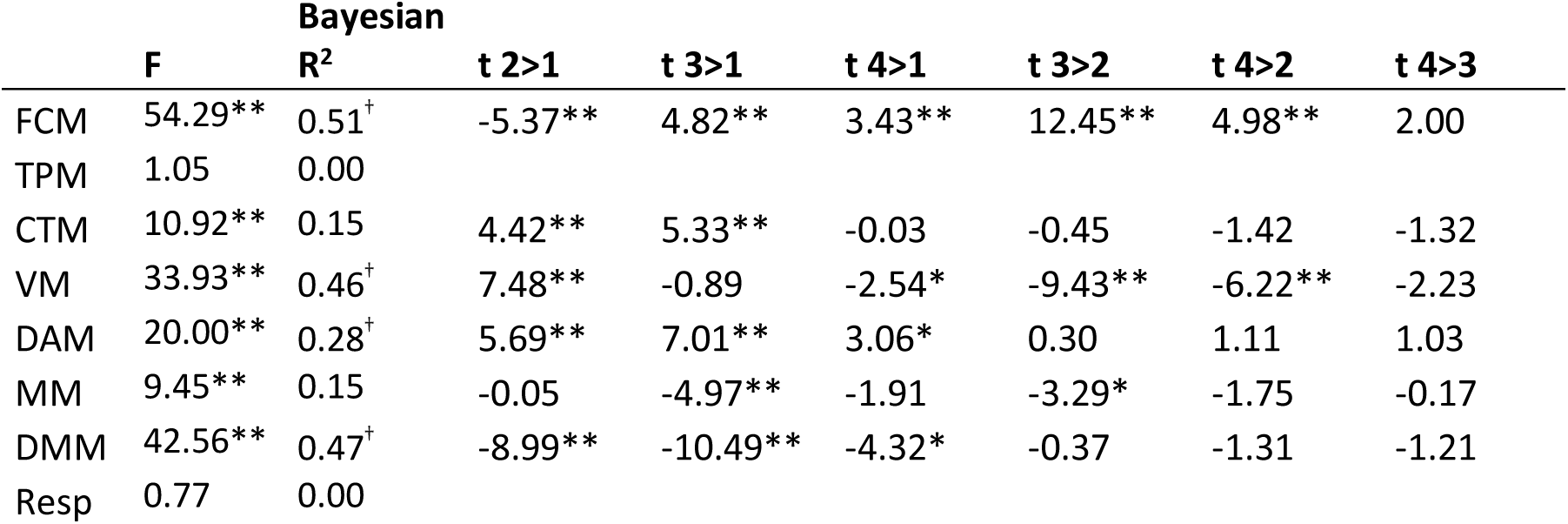
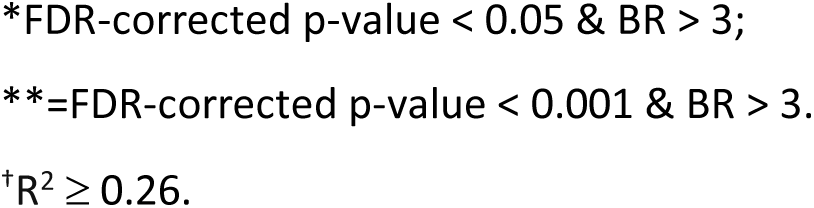
Comparing residual network activity across subgroups.

| | F | Bayesian<br>$R^2$ | t 2>1 | t 3>1 | t 4>1 | t 3>2 | t 4>2 | t 4>3 |
| --- | --- | --- | --- | --- | --- | --- | --- | --- |
| FCM | 54.29** | 0.51<sup>†</sup> | -5.37** | 4.82** | 3.43** | 12.45** | 4.98** | 2.00 |
| TPM | 1.05 | 0.00 |  |  |  |  |  |  |
| CTM | 10.92** | 0.15 | 4.42** | 5.33** | -0.03 | -0.45 | -1.42 | -1.32 |
| VM | 33.93** | 0.46<sup>†</sup> | 7.48** | -0.89 | -2.54* | -9.43** | -6.22** | -2.23 |
| DAM | 20.00** | 0.28<sup>†</sup> | 5.69** | 7.01** | 3.06* | 0.30 | 1.11 | 1.03 |
| MM | 9.45** | 0.15 | -0.05 | -4.97** | -1.91 | -3.29* | -1.75 | -0.17 |
| DMM | 42.56** | 0.47<sup>†</sup> | -8.99** | -10.49** | -4.32* | -0.37 | -1.31 | -1.21 |
| Resp | 0.77 | 0.00 |  |  |  |  |  |  |

\*FDR-corrected p-value < 0.05 & BR > 3; \*\*=FDR-corrected p-value < 0.001 & BR > 3. <sup>†</sup>R<sup>2</sup> ≥ 0.26.

The most substantial differences in residual brain activity were observed for the FCM, VM, DAM, and DMM, as reflected by the corresponding Bayes Factors and values of Bayesian R^2^ (all ≥ 0.26, indicating substantial effect size; Cohen, 2013). Focusing on four most influential modules, subgroup 1 is characterized by less FCM activity compared to subgroups 3 and 4, the lowest DAM activity across all groups, and the least negative DMM activity. Subgroup 2 demonstrates the least FCM activity but the highest VM activity across all subgroups. In contrast, group 3 shows higher FCM activity than subgroups 1 and 2, but lower VM activation than subgroup 2. Subgroup 4 shows a similar pattern as subgroup 3 with even higher FCM and lower VM activity. We observed no significant differences for the TPM and overall responsivity of brain activity to the task load manipulation. These results remained consistent after applying age corrections.

In summary, while subgroup 2 exhibited the lowest frontal activation paired with higher visual activation, subgroups 3 and 4 displayed the opposite pattern. In contrast, subgroup 1 was characterized by having the least DAM activity and DMM deactivation. This may suggest that different neural mechanisms were used to complete the VSTM task, with some individuals relying more heavily on visual processing, others engaging frontal control networks, and or showing weaker engagement of task-related networks overall.

### 3.4 Subgroups showed no differences in demographics or VSTM task performance

Subgroup characteristics are displayed in **Table 2**. No significant between-group differences were found for age, sex, education (all p > 0.05, BF < 3). We found a trend level age difference between the subgroups (F_3,40.8_=2.79, p = 0.053, BF = 0.43), with subgroup 4 tending to be younger than the other participant groups (subgroup 1: t_23.6_ = 2.83, p = 0.009, BF = 3.70; subgroup 2: t_27.7_ = 2.29, p = 0.030, BF = 1.74; subgroup 3: t_18_ = 2.01, p = 0.059, BF = 1.49; **Figure 4**). VSTM performance did not significantly differ across subgroups (precision and number of items in memory at load 3, all p > 0.05, BF < 3). These results were unaffected by age corrections. A complete overview of the between-group statistics for the subgroup characteristics can be found in **Supplementary Table 3**.

**Figure 4:**
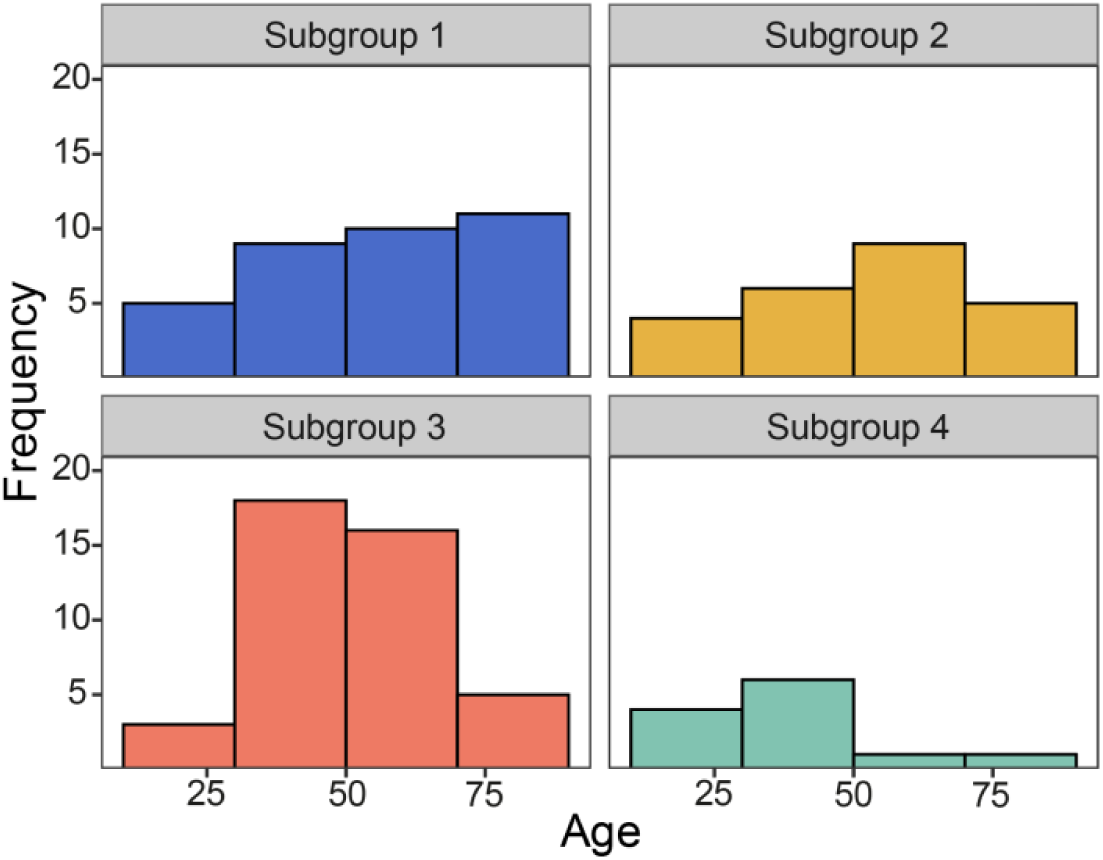
Distribution of age across subgroups.

**Table 2.** Subgroup characteristics.

|  | All participants | Subgroup 1 | Subgroup 2 | Subgroup 3 | Subgroup 4 |
| --- | --- | --- | --- | --- | --- |
| *N* | 113 | 35 | 24 | 42 | 12 |
| Age, mean (SD) | 53 (18.0) | 57 (19.1) | 55 (19.8) | 51 (15.6) | 41 (15.4) |
| Sex |  |  |  |  |  |
| Male, n (%) | 59 (52%) | 14 (40%) | 15 (63%) | 24 (57%) | 6 (50%) |
| Female, n (%) | 54 (48%) | 21 (60%) | 9 (38%) | 18 (43%) | 6 (50%) |
| Highest education |  |  |  |  |  |
| University, n (%) | 82 (73%) | 21 (60%) | 19 (79%) | 32 (76%) | 10 (83%) |
| A' levels, n (%) | 17 (15%) | 6 (17%) | 3 (13%) | 7 (17%) | 1 (8.3%) |
| GCSE grade, n (%) | 9 (8%) | 4 (11%) | 1 (4.2%) | 3 (7.1%) | 1 (8.3%) |
| None > 16, n (%) | 5 (4.4%) | 4 (11%) | 1 (4.2%) | 0 (0%) | 0 (0%) |
| Number of items in memory (load 3), mean (SD) | 2.40 (0.46) | 2.35 (0.50) | 2.38 (0.47) | 2.43 (0.47) | 2.49 (0.34) |
| Memory precision (load 3), mean (SD) | 0.12 (0.05) | 0.10 (0.04) | 0.12 (0.04) | 0.12 (0.05) | 0.13 (0.04) |

### 3.5 Subgroups revealed distinct associations between brain activity and VSTM task performance

Next, we investigated the association between brain activity and task performance across all participants, and separately within each subgroup, while correcting for age. From these analyses, we excluded subgroup 4 because of its small number of participants (*n* = 12, yielding only 16% power to detect a medium effect [r = 0.30] at alpha = 0.05, two-sided). To limit the number of comparisons, we focused on the brain modules with the most pronounced differences between the subgroups: the FCM, VM, DAM, and DMM. A complete overview of the associations between brain module activity and performance on the VSTM task for each of the subgroups is displayed in **Supplementary Table 4.**

For the FCM, analyses across all participants revealed no significant associations between FCM activity and memory precision (r = −0.15, 95% CI = [−0.33; 0.05], uncorrected p = 0.136, BF = 0.45; **Figure 5A**). However, looking into the separate subgroups, we found that subgroups 1 and 2 showed negative associations between FCM activity and memory precision (subgroup 1: r = −0.37, 95% CI = [−0.62; −0.04], uncorrected p = 0.030, BF = 2.36; subgroup 2: r = −0.41, 95% CI = [−0.70; 0.01], uncorrected p = 0.045, BF = 1.93, see **Figure 5B**), albeit not surviving corrections for multiple comparisons (FDR-adjusted p = 0.059, and p = 0.180, respectively). This pattern was absent in subgroup 3 (r = 0.12, p = 0.465, BF = 0.30). When we compared the slopes between subgroups 1 and 3, we found that these tended to be different (z = −2.11, 95% CI = [−0.87; −0.03], uncorrected p = 0.035, FDR-adjusted p = 0.060), the same was true for the comparison between subgroups 2 and 3 (z = −2.05, 95% CI = [−0.94; −0.02], uncorrected p = 0.040, FDR-adjusted p = 0.060). These findings illustrate how certain associations between brain activity and performance may only become apparent at the subgroup level. Notably, subgroups 1 and 2 were characterized by having relatively low FCM recruitment compared to subgroups 3 and 4. Therefore, such negative associations suggest that the low recruitment of the FCM within subgroups 1 and 2 may have been beneficial to their memory precision. No association was observed between FCM activity and number of items in memory in any of the groups (all p > 0.05).

**Figure 5:**
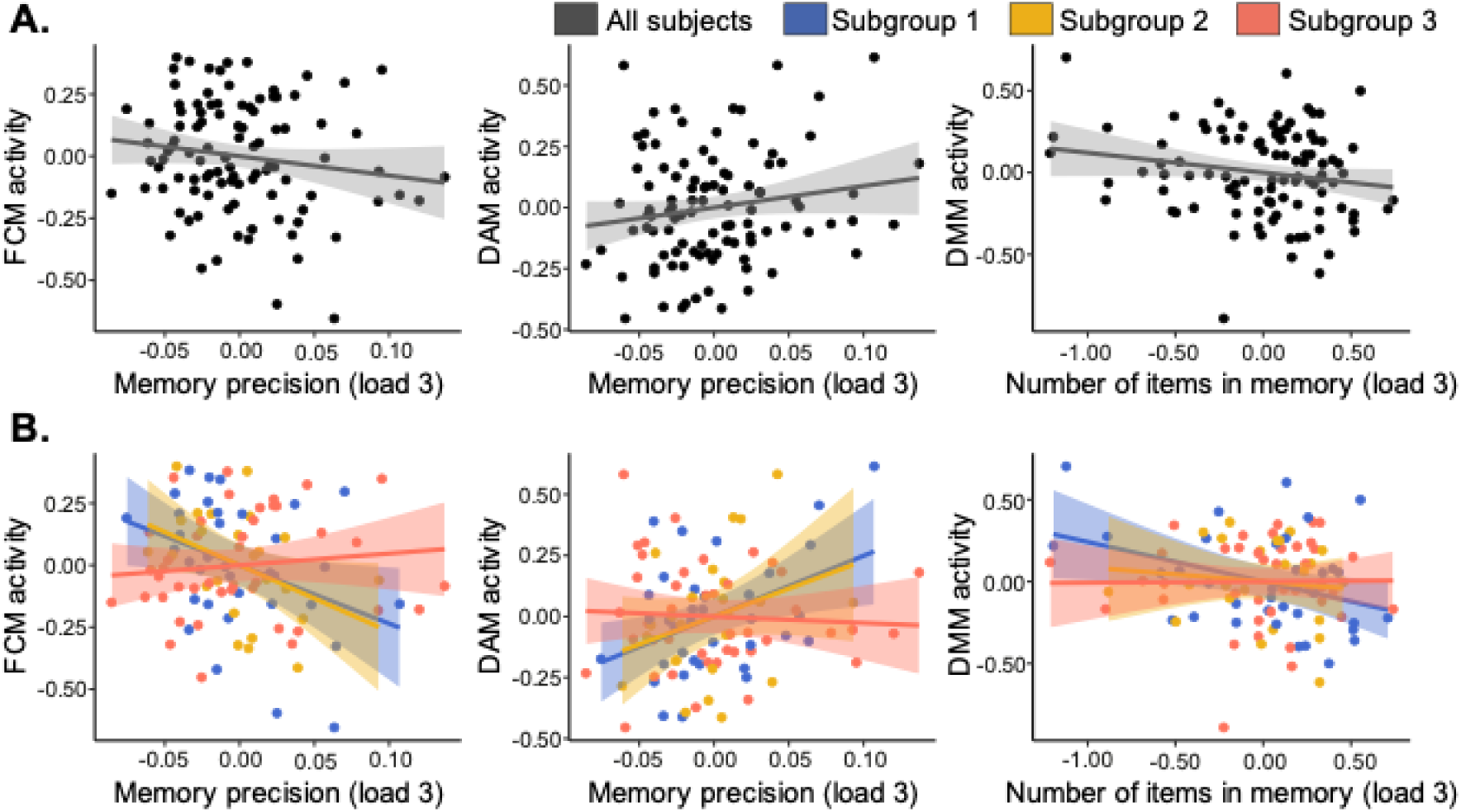
Associations between brain module activity in the frontal control module (FCM), dorsal attention module (DAM), and default mode module (DMM) and performance on the visual short-term memory task. All analyses were corrected for age. **A**. Whole-group analyses revealed no associations between brain module activity and performance. **B.** Subgroup specific associations between brain module activity and performance.

For the VM, no significant associations with items in memory or memory precision were observed (all p > 0.05).

In the DAM, we did not observe significant associations between DAM activity and memory precision across all participants (r = 0.17, 95% CI = [-0.03; 0.35], uncorrected p = 0.098, BF = 0.58). However, when considering each subgroup separately, we observed that subgroup 1 demonstrated a positive association between DAM activity and memory precision (r = 0.42, 95% CI = [0.10; 0.66], FDR-adjusted p = 0.048, BF = 4.97, see **Figure 5B).** In subgroup 2, this association tended in a similar direction; however, this was not significant (r = 0.30, uncorrected p = 0.152), whereas this association was absent for subgroup 3 (r = −0.06, uncorrected p = 0.69). The slope in subgroup 1 differed from that in subgroup 3 (z = 2.14, 95% CI = [0.04; 0.86], uncorrected p = 0.032) but did not survive FDR-corrections (FDR-adjusted p = 0.097). Subgroup 1 was characterized by having the least DAM recruitment. This suggests that for subgroup 1, their low DAM activity may have negatively impacted their memory precision. There were no significant associations between DAM activity and number of items in memory in any of the groups.

In the DMM, no associations were observed between DMM activity and memory precision across the whole group or subgroups (all p > 0.05). For the number of items in memory, no significant associations were observed in the whole group (r = −0.18, 95% CI = [-0.36; 0.02], uncorrected p = 0.073, BF = 0.74). Within subgroup 1 a negative association was observed between DMM deactivation and the number of items in memory (r = −0.38, 95% CI = [−0.63; - 0.05], uncorrected p = 0.026, BF = 2.65), suggesting that the increased activity (or reduced suppression) of the DMM in subgroup 1 may have hindered the number of items participants in that subgroup could recall. However, this association did not survive FDR-corrections (FDR-adjusted p = 0.102). No associations were found in subgroup 2 (r = −0.11, p = 0.597, BF = 0.35) or subgroup 3 (r = 0.01, p = 0.942, BF = 0.23; see **Figure 5B**). The slopes did not significantly differ between groups.

Taken together, these results illustrate how associations between brain module activity and VSTM performance can be subgroup-specific, even though task performance did not differ across groups. This further supports the notion that these distinct brain activity profiles reflect different yet equally effective pathways to perform the task. Additionally, these findings illustrate how whole-group analyses may mask meaningful brain-behaviour relationships that only emerge at the subgroup level.

### 3.6 Subgroups differ in white matter integrity

Next, we investigated whether structural brain characteristics differed between groups. First, we compared grey matter volumes within each of the modules between groups while correcting for total intracranial volume. Grey matter volume in the TPM significantly differed across subgroups (F_3,39.84_ = 5.67, FDR-adjusted p-value = 0.017, BF = 5.10), however this effect disappeared after additionally correcting for age (unadjusted p = 0.048; BF = 0.72).

We also investigated whether the different subgroups showed significant differences in white matter microstructure, as measured by mean kurtosis (MK). We found significant differences between subgroups for four white matter tracts (**see Figure 6**). These included the left inferior frontal occipital fasciculus (IFOF; F_3,37.8_ = 4.73, FDR-adjusted p = 0.030, BF = 5.04), right IFOF (F_3,36.10_ = 4.08, FDR-adjusted p-value = 0.049, BF = 3.49), left inferior longitudinal fasciculus (F_3,35.09_ = 6.71, FDR-adjusted p-value = 0.011, BF = 28.10) and the left uncinate fasciculus (F_3,35.43_ = 6.60, FDR-adjusted p-value = 0.011, BF = 16.93).

**Figure 6:**
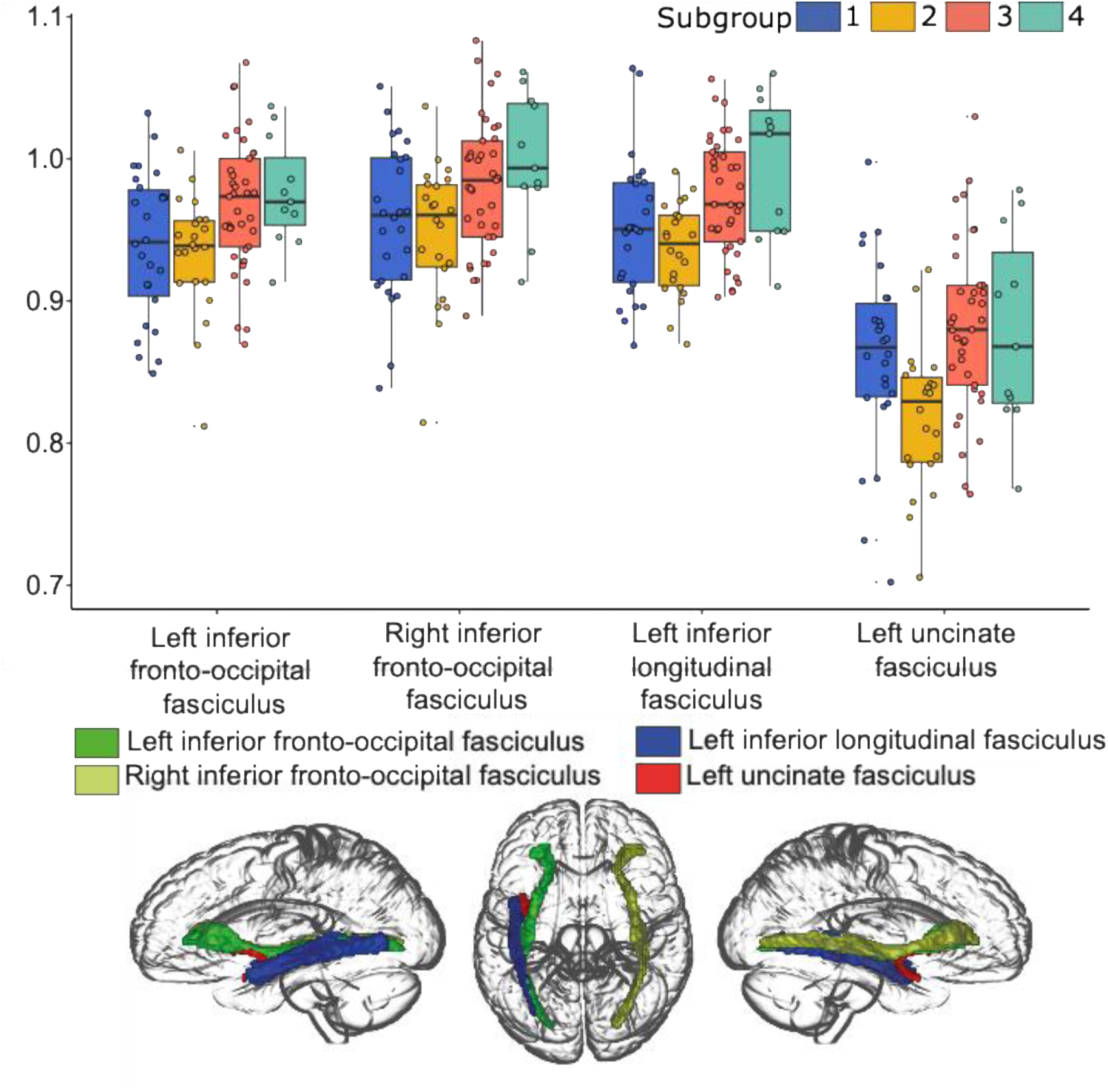
The upper part of the figure shows the average mean kurtosis values in each subgroup for the white matter tracts that displayed significant between-group differences (FDR-corrected p-value < 0.05 and Bayes Factor > 3). The bottom part visualizes the white matter tracts in a glass brain, generated with MRIcroGL (https://www.nitrc.org/projects/mricrogl). Only the effects in the left inferior longitudinal fasciculus and left uncinate fasciculus survived additional age corrections.

Only the effects in the left inferior longitudinal fasciculus and the left uncinate fasciculus remained significant after correcting for age (F_3,34.8_ = 5.42, FDR-adjusted p-value = 0.033, BF = 5.81; F_3,35.4_ = 6.64, FDR-corrected p-value = 0.020, BF = 17.32, respectively). The left inferior fasciculus showed lower MK in subgroup 2 relatively to subgroup 3 (t_53.56_ = 3.71, FDR-corrected p = 0.003, BF = 30.24) and subgroup 4 (t_13.37_ = 2.65, unadjusted p = 0.019, BF = 11.23), however the latter effect did not survive FDR-corrections.

The left uncinate fasciculus showed lower mean kurtosis in subgroup 2 relatively to subgroup 1 (t_45.50_ = 2.58, FDR-corrected p-value = 0.039, BF = 3.49), subgroup 3 (t_50.46_ = 4.37, FDR-corrected p-value < 0.001, BF > 100), and subgroup 4 (t_15.49_ = 2.60, FDR-corrected p-value = 0.039, BF = 6.62). So, subgroup 2, which is the subgroup with the least FCM activity but the highest VM activity across all subgroups, specifically showed lower MK in the left uncinate fasciculus and the left inferior longitudinal fasciculus. Notably, the low recruitment of the FCM within this subgroup appeared to be beneficial to their memory precision.

A complete overview of the between-group statistics for grey matter volumes and MK is provided in **Supplementary Tables 4 and 5.**

## 4 Discussion

While we often assume that similar patterns of neural activity are engaged to complete a cognitive task across individuals, accumulating evidence suggests that this is not necessarily the case, even in the absence of brain injury or disease (Noppeney et al., 2004; Price & Friston, 2002; Seghier & Price, 2018). Instead of discarding inter-individual variability in task-related functional activity, we leveraged it in our analyses to investigate brain degeneracy or many-to-one mapping in the context of VSTM performance. We identified four participant subgroups with distinct brain activity profiles, particularly within the FCM, VM, DAM, and DMM. Despite substantial differences in brain activity, these subgroups showed no significant differences in task performance or age, suggesting that these profiles may reflect different yet equally effective pathways to perform the task. This was further supported by the small, albeit distinct associations between task performance and brain modular activity across subgroups. Individual differences in brain structure may partly account for differences in brain activity, as subgroups showed age-independent differences in white matter integrity for the left inferior longitudinal fasciculus and the left uncinate fasciculus. Here, we first discuss our findings in relation to many-to-one mapping. Subsequently, we describe the modules and subgroups that we identified in more detail as well as the role of global versus local inter-individual variability in brain activity. Finally, we highlight the factors that contributed to subgroup variations in brain activity. Given the exploratory nature of our analyses, we also consider how the observed subgroup differences can serve as a starting point for testable hypotheses, guiding future research on individual differences in brain function.

Degeneracy suggests that there can be multiple brain activation patterns leading to the same behavioural outcome (Edelman & Gally, 2001; Mason et al., 2015; Price & Friston, 2002; Westlin et al., 2023). We identified four subgroups with marked differences in recruitment of various brain modules during the VSTM task, after accounting for differences in overall network activity. These results add to previous studies that revealed subgroups with distinct profiles of task-related functional activity (Cerliani et al., 2017; Fischer-Baum et al., 2018; Kherif et al., 2009; Noppeney et al., 2006; Noppeney et al., 2005; Seghier et al., 2008; Seghier & Price, 2016). Similarly to previous findings (Fischer-Baum et al., 2018; Noppeney et al., 2006), these subgroups demonstrated similar levels of performance. These findings illustrate how different patterns of brain activity can lead to the same functional outcome, providing support for the notion of brain degeneracy. Additionally, we observed several unique associations between brain modular activity and task performance. This finding suggests that there is not a single route to successful cognitive performance, and the optimal route may vary from person to person. This finding highlights the individual-specific nature of brain-cognition relationships, where diverse neural mechanisms can yield similar outcomes (Noppeney et al., 2006; Seghier & Price, 2018).

As a first step toward identifying subgroups, we reduced the dimensionality of our data, identifying seven brain modules that showed similar between-participant variability in their activity patterns during the maintenance period of the VSTM task (load 3 vs. load 1). Strikingly, these modules show strong correspondence to brain networks that are typically identified using functional connectivity analyses, such as the frontoparietal control network and default mode network (Power et al., 2011; Yeo et al., 2011). This happens even though functional connectivity is typically based on temporal correlations in activity within participants while we investigated correlations in activity between participants, suggesting that there is a shared underlying source. These results corroborate the idea that there is a stable intrinsic network architecture that is largely uniform across individuals but also shows reliable individual differences (Finn et al., 2015; Fox & Raichle, 2007; Gratton et al., 2018). This intrinsic architecture manifests itself both in how networks are functionally connected within a single person (within-subject) and in the differences in network activity among individuals (between-subjects). In line with this, previous studies demonstrated that functional connectivity can be used to predict task-activation, and that individual differences in connectivity are related to variability in task-related activation (Cole et al., 2014; Cole et al., 2021; Finn et al., 2015; Tavor et al., 2016). From a theoretical point of view, it has been proposed that intrinsic functional connectivity serves as a large-scale analogue of synaptic weight structure (Mill et al., 2022; Wig et al., 2011), providing a mechanistic account of how intrinsic network architecture may constrain individual patterns of brain activity.

We observed that the brain activity across the seven brain modules was highly correlated, corroborating previous findings where the global recruitment of task-related functional activity explained most inter-individual variability, both in relation to performance and age (Chen et al., 2022; Mitchell et al., 2023; Samu et al., 2017). Consistent with prior studies, the average activity across all modules explained 60-75% of the variability in activity observed within each brain module. To be able to look beyond this general factor, we used module-specific activity for our subsequent analysis, where the average activity within each module was corrected for the average activity across all modules. Results from the latent profile analysis suggests that there is meaningful inter-individual variability beyond this general factor, by revealing subgroups with differences in white matter integrity and unique associations between brain activity and performance. These findings illustrate how individual differences in task-related brain activation arise from both overall responsivity to the task (Samu et al., 2017), and the specific recruitment patterns of brain regions, highlighting the importance of individual-specific factors in shaping task-related brain activity (Cole et al., 2021; Finn et al., 2015; Gratton et al., 2018).

Of the seven modules we identified, four showed the most significant differences in activity across subgroups: the FCM, VM, DAM, and DMM. These brain modules largely overlap with the frontoparietal network, visual network, dorsal attention network, and default mode network, respectively (Power et al., 2011; Yeo et al., 2011). With increasing task load, activity typically increases in the frontoparietal and dorsal attention networks, whereas the default mode network becomes more suppressed (Huang et al., 2016; Nagel et al., 2009; Salami et al., 2019; Salami et al., 2018; Zuo et al., 2019). This reflects a shift away from internally focused processes to task-oriented activity (Fox et al., 2005; Seeley et al., 2007; Smith et al., 2009). Notably, this pattern was weaker for subgroup 1, suggesting that this subgroup might have been less attentive during the maintenance phase (Anticevic et al., 2012; Berry et al., 2017). Surprisingly, this subgroup showed task performance comparable to the other subgroups, despite the positive association between DAM activity and memory precision, and the negative association between DMM activity and the number of items in memory. One possibility is that VSTM performance was sustained in this subgroup through increased engagement during the encoding or retrieval phase (Linke et al., 2011; Magen, 2017). Stimulus information might also have been preserved in a passive state, where maintenance is facilitated by neural mechanisms not characterized by increased activity (i.e., activity-silent mechanisms), such as short-term synaptic plasticity (Lewis-Peacock et al., 2014; Stokes, 2015).

Successful VSTM performance is further facilitated through interactions between the frontoparietal and visual networks (D’Esposito & Postle, 2015; Eriksson et al., 2015; Scimeca et al., 2018). The sensory recruitment hypothesis of working memory suggests that the same brain regions responsible for encoding the stimulus are also essential for maintaining this information (D’Esposito, 2007; D’Esposito & Postle, 2015). In line with this, in the context of VSTM, visual regions have been shown to maintain detailed sensory representations of the stimuli (Emrich et al., 2013; Postle, 2015; Serences, 2016). The frontoparietal network is thought to exert top-down control over these areas and support the maintenance of more abstract representations of the stimuli, such as verbal descriptions (Christophel et al., 2017; Gayet et al., 2018; Scimeca et al., 2018; Serences, 2016).

Interestingly, subgroup 2 demonstrated the highest VM activity and lowest FCM activity. The reduced FCM recruitment in this subgroup was beneficial to VSTM performance, as reflected by the negative associations between FCM activity and memory precision in these participants. In contrast, subgroups 3 and 4 demonstrated relatively high FCM activity and lower VM activity. These results provide an interesting starting point for several hypotheses. The use of different strategies to complete the task is an important source of variability for task-related brain activity (Miller et al., 2012; Sanfratello et al., 2014). Consistently, previous studies that identified subgroups with distinct brain activity profiles during task engagement attributed this variability to the adoption of different strategies (Kherif et al., 2009). It is possible that subgroup 2 relied more on visual representations, while subgroup 3 and 4 maintained stimulus information through more abstract forms, thereby relying more on the FCM. This would also align with the two most commonly reported strategies to perform VSTM tasks: maintaining visual representations of the stimuli (e.g., picturing the red dots moving to the left corner) and transforming salient stimulus features into verbal representations (e.g., mentally repeating “the red dots moved to the left corner”) (Berger & Gaunitz, 1979; Pearson & Keogh, 2019). Therefore, one hypothesis is that subgroup differences in neural activity reflect distinct cognitive strategies, with VM recruitment supporting visual imagery and FCM recruitment reflecting verbal or abstract strategies.

Subgroup 2 also showed decreased white matter integrity in the inferior longitudinal fasciculus and the uncinate fasciculus, whereas this was preserved in subgroups 3 and 4. The inferior longitudinal fasciculus is involved in modulating visual information and semantic-lexical processing (Herbet et al., 2018; Shekari & Nozari, 2023). The uncinate fasciculus is associated with associative learning and semantic retrieval (Olson et al., 2015; Von Der Heide et al., 2013). This suggests that the use of a visually oriented strategy in subgroup 2 may be linked to the reduced microstructural integrity in two white matter tracts that play a role in creating and retrieving verbal representations (Duffau et al., 2013; Herbet et al., 2018; Shekari & Nozari, 2023). This pattern leads us to another hypothesis: structural constraints, such as reduced integrity in tracts supporting the formation of abstract representations, contribute to the use of alternative strategies. However, we did not find any direct association between brain activity and performance in subgroup 3, which makes it challenging to interpret how this brain activity profile precisely facilitated task performance in this group.

Several methods can be employed to test these hypotheses in future studies. Self-reported questionnaires or semi-structured interviews could be used (e.g., Miller et al., 2012). Reported strategies could also inform alternative clustering approaches (e.g., Sanfratello et al., 2014), allowing investigation of whether subgroups derived from strategy reports resemble those identified from neural data. Furthermore, experimental manipulations of strategy use, such as instructing participants to rely either on visual representations or verbal representations (e.g., Kraemer et al., 2017), would be expected to alter the balance between VM and FCM engagement. Concurrent measurement of structural brain correlates in this context would enable testing of whether individual variation in white matter integrity biases participants toward certain cognitive strategies. Lesion-symptom mapping provides a complementary approach in patient populations (e.g., Lugtmeijer et al., 2021), to test whether localized brain damage increases reliance on alternative strategies. Individuals with aphantasia, a condition characterized by the absence of visual imagery but preserved visual working memory (Pearson & Keogh, 2019), also provide an interesting comparison group. As these individuals rely on non-visual strategies to retain comparable performance (Keogh et al., 2021; Zeman, 2024), contrasting their brain activation patterns and structural brain correlates with controls may help disentangle visual from non-visual routes to successful task performance.

The age-independent differences in white matter integrity across subgroups were consistent with our expectation that structural brain characteristics would contribute to the variability in brain activity among the subgroups (or vice versa). This also aligns with previous research linking white matter microstructure to the recruitment of different brain regions during task performance (Burzynska et al., 2013; Burzynska et al., 2015). We did not observe any differences in grey matter across subgroups, suggesting that white matter integrity may be relatively more important to explain inter-individual variability in brain activity in our sample (Miller et al., 2012).

Typically, white matter degradation is associated with declines in cognitive performance, including working memory (Bennett & Madden, 2014; Oschwald et al., 2020). Our results suggest that this is not necessarily the case when different brain activation patterns can be used to perform a task (Madden et al., 2009), corresponding to the notion of brain degeneracy (Noppeney et al., 2004; Price & Friston, 2002). It should be noted that the differences in white matter integrity between subgroups were relatively small, emphasizing the need to further investigate these associations in future studies. Nevertheless, our findings underscore that individual differences in structural brain characteristics are an important source of variability in brain activity and cognitive performance (Genon et al., 2022; Kanai & Rees, 2011; Miller et al., 2012; Seghier & Price, 2018).

Although prior research has demonstrated strong age-related differences in brain activity and performance during VSTM tasks, including the one used here (Lugtmeijer et al., 2023; Morcom & Henson, 2018; Sander et al., 2012), we did not observe any significant differences in age or performance between the subgroups. This suggests that other factors beyond age or task performance were the primary drivers behind the distinct brain activation patterns observed across subgroups (Finn et al., 2015; Gratton et al., 2018). These factors may have included the extent to which participants were engaged throughout the task (Linke et al., 2011; Magen, 2017; Rose, 2020), the strategies adopted to complete the task (Kherif et al., 2009; Miller et al., 2012), and variations in white matter integrity (Burzynska et al., 2013; Burzynska et al., 2015), as discussed above. Nonetheless, age was positively correlated with the global recruitment of brain activity, consistent with prior research showing that that variability in overall responsivity was primarily due to age-differences (Mitchell et al., 2023; Samu et al., 2017). Therefore, the lack of age differences between subgroups might also be explained by our approach of clustering participants based on the residual brain activity in each module (i.e., after regressing out the overall responsiveness). In contrast, task performance was not associated with overall responsivity, suggesting that this factor is not the most substantial driver of inter-individual variability in brain activity. Therefore, studies that are interested in explaining performance metrics would benefit from distinguishing individuals based on performance itself rather than brain activity (e.g., Lövden et al., 2018; Salami et al., 2018).

There are several limitations that should be considered. First, our study relied on several data-driven methods, including the Louvain modularity algorithm and LPA to facilitate the identification of hidden subgroups of individuals with different brain activity patterns during the VSTM task. While these approaches are valuable to uncover novel patterns that might be overlooked by traditional univariate methods, they do have limitations. The Louvain algorithm is stochastic in nature and may yield different results across iterations, and it constrains each ROI to a single module, even though regions may participate in multiple functional networks (Lancichinetti & Fortunato, 2012; Rubinov & Sporns, 2011; Traag et al., 2019). Furthermore, the strong correspondence between our study-specific modules and conventional functional networks suggests that the use of study-specific modules is not a crucial part of our pipeline, and the use of predefined networks would likely yield similar results. The use of LPA, while flexible, is sensitive to model specification and sample characteristics, and may overfit or yield spurious classes in small samples (Scrucca et al., 2016; Spurk et al., 2020). As with all data-driven methods, results should be interpreted with caution and validated in independent datasets.

A second limitation concerns the study design. The VSTM task design used fixed durations without temporal jitter between encoding, maintenance, and response phases. Given the extended nature of the BOLD response, this makes it difficult to fully dissociate these closely spaced phases. Although prior work using this paradigm showed that the observed load-modulated effects were not fully attributable to spillover artefacts, a complete dissociation remains impossible (cf. Lugtmeijer et al., 2023). Moreover, we merely focused on brain-behaviour associations. While this approach is useful to characterize the various factors underlying subgroup differences in brain activity, brain-behaviour prediction approaches – where models are evaluated in held-out data – are more robust to noise and offer clearer translational potential (Mill et al., 2020; Poldrack et al., 2020). It should also be noted that the use of a single-task approach limits the generalizability of findings compared to multi-task paradigms (King et al., 2019; Mill & Cole, 2023).

Finally, the interpretation of subgroup differences should be made with caution. To interpret our findings, we explored how the brain activity differences across subgroups may be associated with different cognitive processes. Although the use of different strategies has been recognized an important source of task-related functional activity (Kherif et al., 2009; Miller et al., 2012; Noppeney et al., 2006; Sanfratello et al., 2014), we note that this interpretation remains speculative as we do not have any information about which strategies participants used when they performed the task. Therefore, the evidence here should be taken as proof of principle, showing that when we use a different approach to look at task-related brain activity, interesting dimensions of inter-individual variability are revealed that may otherwise remain hidden (Cerliani et al., 2017; Fischer-Baum et al., 2018; Kherif et al., 2009; Noppeney et al., 2006; Seghier et al., 2008; Seghier & Price, 2016; Seghier & Price, 2018). However, as illustrated in this discussion, our results can be used as a starting point for testable hypotheses in future studies. Furthermore, our dataset consists of a relatively small number of participants, which may have limited our ability to observe within-group associations with performance. Although we excluded subgroup 4 (n = 12) from these analyses, the remaining subgroup sizes (n = 35, n = 24, n = 42) correspond to 30-50% power to detect a medium effect (r = 0.30) and 73–93% power for a large effect (r = 0.50) at α = 0.05 (two-sided). Accordingly, these findings should be interpreted cautiously and emphasize the need for replication with larger sample sizes, preferably > 80 participants per subgroup to achieve 80% power for medium effects. Lastly, although we discussed the role of several factors in explaining the variability in brain activity between subgroups, this does not rule out the possibility that other variables that were not considered here could also contribute, such as other early-life factors, genetic factors or brain development (Walhovd et al., 2023). We hope that future studies focusing on this subject will include larger samples and a more extensive characterization of individual-specific factors that could further explain this variability.

Taken together, the novel approach introduced here allowed us to uncover distinct subgroups with marked differences in neural activity during the VSTM task, associations between brain module activity and VSTM performance, and white matter microstructural properties. However, these subgroups did not significantly differ in their demographic characteristics or VSTM performance. The identification of distinct subgroups illustrates how multiple brain activation patterns may underlie successful VSTM performance. This further highlights the importance of considering brain degeneracy to understand variability in brain-cognition associations (Edelman & Gally, 2001; Mason et al., 2015; Price & Friston, 2002; Westlin et al., 2023). Recognizing this possibility provides us with a more comprehensive understanding of the different ways in which our brains support cognitive functions and how this may vary from person to person. Consequently, these insights may help to unravel new ways in which structure and function interact, and how complementary routes can lead to the same cognitive outcomes.

## Data and code availability

Data availability is provided via the Cam-CAN data portal (https://camcan-archive.mrc-cbu.cam.ac.uk/dataaccess/). The scripts and analysis pipeline are made available via Github (https://github.com/michellegjansen/neural_pathways_VSTM).

## Author contributions

Michelle G. Jansen: Conceptualization, methodology, software, investigation, visualization, validation, data curation, formal analysis, writing – original draft. Alireza Salami: Conceptualization, methodology, writing – review & editing. Fernando-Ruiz Martínez: Methodology, software, investigation, data curation, formal analysis. Daniel J. Mitchell: Methodology, data curation, formal analysis, writing – review & editing. Joukje M. Oosterman: Supervision, writing – review & editing. Linda Geerligs: Conceptualization, methodology, software, investigation, visualization, validation, data curation, formal analysis, supervision, writing – original draft.

## Acknowledgments

We thank the Cam-CAN respondents and their primary care teams in Cambridge for their participation in this study. The Cam-CAN team involves many different researcher and support staff, who made this study possible. We would like to express our gratitude to the project principal personnel, research associates, research assistants, affiliated personnel, research interviewers, and administrative staff. Further information about the Cam-CAN corporate authorship membership can be found at https://cam-can.mrc-cbu.cam.ac.uk/corpauth/#16.

## References

Anticevic, A., Cole, M. W., Murray, J. D., Corlett, P. R., Wang, X. J., & Krystal, J. H. (2012). The role of default network deactivation in cognition and disease. Trends Cogn Sci, 16(12), 584–592. 10.1016/j.tics.2012.10.008

Ashburner, J., & Friston, K. J. (2005). Unified segmentation. Neuroimage, 26(3), 839–851. 10.1016/j.neuroimage.2005.02.018

Bassett, D. S., Porter, M. A., Wymbs, N. F., Grafton, S. T., Carlson, J. M., & Mucha, P. J. (2013). Robust detection of dynamic community structure in networks. Chaos, 23(1), 013142. 10.1063/1.4790830

Bays, P. M., Catalao, R. F., & Husain, M. (2009). The precision of visual working memory is set by allocation of a shared resource. Journal of vision, 9(10), 7–7.

Benitez, A., Jensen, J. H., Falangola, M. F., Nietert, P. J., & Helpern, J. A. (2018). Modeling white matter tract integrity in aging with diffusional kurtosis imaging. Neurobiology of Aging, 70, 265–275. 10.1016/j.neurobiolaging.2018.07.006

Benjamini, Y., & Hochberg, Y. (1995). Controlling the False Discovery Rate: A Practical and Powerful Approach to Multiple Testing. Journal of the Royal Statistical Society: Series B (Methodological), 57(1), 289–300. 10.1111/j.2517-6161.1995.tb02031.x

Bennett, I. J., & Madden, D. J. (2014). Disconnected aging: cerebral white matter integrity and age-related differences in cognition. Neuroscience, 276, 187–205. 10.1016/j.neuroscience.2013.11.026

Berger, G. H., & Gaunitz, S. C. B. (1979). Self-rated imagery and encoding strategies in visual memory. British Journal of Psychology, 70(1), 21–24. 10.1111/j.2044-8295.1979.tb02137.x

Berry, A. S., Sarter, M., & Lustig, C. (2017). Distinct Frontoparietal Networks Underlying Attentional Effort and Cognitive Control. J Cogn Neurosci, 29(7), 1212–1225. 10.1162/jocn_a_01112

Buades-Rotger, M., Smeijers, D., Gallardo-Pujol, D., Krämer, U. M., & Brazil, I. A. (2023). Aggressive and psychopathic traits are linked to the acquisition of stable but imprecise hostile expectations. Translational Psychiatry, 13(1), 197. 10.1038/s41398-023-02497-0

Burzynska, A. Z., Garrett, D. D., Preuschhof, C., Nagel, I. E., Li, S.-C., Bäckman, L., Heekeren, H. R., & Lindenberger, U. (2013). A Scaffold for Efficiency in the Human Brain. The Journal of Neuroscience, 33(43), 17150–17159. 10.1523/jneurosci.1426-13.2013

Burzynska, A. Z., Wong, C. N., Voss, M. W., Cooke, G. E., McAuley, E., & Kramer, A. F. (2015). White Matter Integrity Supports BOLD Signal Variability and Cognitive Performance in the Aging Human Brain. PLOS ONE, 10(4), e0120315. 10.1371/journal.pone.0120315

Cabeza, R., Albert, M., Belleville, S., Craik, F. I. M., Duarte, A., Grady, C. L., Lindenberger, U., Nyberg, L., Park, D. C., Reuter-Lorenz, P. A., Rugg, M. D., Steffener, J., & Rajah, M. N. (2018). Maintenance, reserve and compensation: the cognitive neuroscience of healthy ageing. Nat Rev Neurosci, 19(11), 701–710. 10.1038/s41583-018-0068-2

Cabeza, R., Anderson, N. D., Locantore, J. K., & McIntosh, A. R. (2002). Aging Gracefully: Compensatory Brain Activity in High-Performing Older Adults. Neuroimage, 17(3), 1394–1402. 10.1006/nimg.2002.1280

Cerliani, L., Thomas, R. M., Aquino, D., Contarino, V., & Bizzi, A. (2017). Disentangling subgroups of participants recruiting shared as well as different brain regions for the execution of the verb generation task: A data-driven fMRI study. Cortex, 86, 247–259. 10.1016/j.cortex.2016.11.017

Chen, X., Rundle, M. M., Kennedy, K. M., Moore, W., & Park, D. C. (2022). Functional activation features of memory in successful agers across the adult lifespan. Neuroimage, 257, 119276. 10.1016/j.neuroimage.2022.119276

Christophel, T. B., Klink, P. C., Spitzer, B., Roelfsema, P. R., & Haynes, J.-D. (2017). The Distributed Nature of Working Memory. Trends in Cognitive Sciences, 21(2), 111–124. 10.1016/j.tics.2016.12.007

Cohen, J. (2013). Statistical power analysis for the behavioral sciences. Academic press.

Cole, M. W., Bassett, D. S., Power, J. D., Braver, T. S., & Petersen, S. E. (2014). Intrinsic and task-evoked network architectures of the human brain. Neuron, 83(1), 238–251. 10.1016/j.neuron.2014.05.014

Cole, M. W., Ito, T., Cocuzza, C., & Sanchez-Romero, R. (2021). The Functional Relevance of Task-State Functional Connectivity. J Neurosci, 41(12), 2684–2702. 10.1523/jneurosci.1713-20.2021

Cusack, R., Vicente-Grabovetsky, A., Mitchell, D. J., Wild, C. J., Auer, T., Linke, A. C., & Peelle, J. E. (2015). Automatic analysis (aa): efficient neuroimaging workflows and parallel processing using Matlab and XML [Technology Report]. Frontiers in Neuroinformatics, 8. 10.3389/fninf.2014.00090

D’Esposito, M. (2007). From cognitive to neural models of working memory. Philosophical Transactions of the Royal Society B: Biological Sciences, 362(1481), 761–772. doi:10.1098/rstb.2007.2086

D’Esposito, M., & Postle, B. R. (2015). The Cognitive Neuroscience of Working Memory. Annual Review of Psychology, 66(Volume 66, 2015), 115-142. 10.1146/annurev-psych-010814-015031

Driessen, J. M. A., Fanti, K. A., Glennon, J. C., Neumann, C. S., Baskin-Sommers, A. R., & Brazil, I. A. (2018). A comparison of latent profiles in antisocial male offenders. Journal of Criminal Justice, 57, 47–55. 10.1016/j.jcrimjus.2018.04.001

Duffau, H., Herbet, G., & Moritz-Gasser, S. (2013). Toward a pluri-component, multimodal, and dynamic organization of the ventral semantic stream in humans: lessons from stimulation mapping in awake patients [Opinion]. Frontiers in Systems Neuroscience, 7. 10.3389/fnsys.2013.00044

Edelman, G. M., & Gally, J. A. (2001). Degeneracy and complexity in biological systems. Proceedings of the National Academy of Sciences, 98(24), 13763–13768.

Emrich, S. M., Riggall, A. C., LaRocque, J. J., & Postle, B. R. (2013). Distributed Patterns of Activity in Sensory Cortex Reflect the Precision of Multiple Items Maintained in Visual Short-Term Memory. The Journal of Neuroscience, 33(15), 6516–6523. 10.1523/jneurosci.5732-12.2013

Eriksson, J., Vogel, Edward K., Lansner, A., Bergström, F., & Nyberg, L. (2015). Neurocognitive Architecture of Working Memory. Neuron, 88(1), 33–46. 10.1016/j.neuron.2015.09.020

Finn, E. S., Poldrack, R. A., & Shine, J. M. (2023). Functional neuroimaging as a catalyst for integrated neuroscience. Nature, 623(7986), 263–273. 10.1038/s41586-023-06670-9

Finn, E. S., Shen, X., Scheinost, D., Rosenberg, M. D., Huang, J., Chun, M. M., Papademetris, X., & Constable, R. T. (2015). Functional connectome fingerprinting: identifying individuals using patterns of brain connectivity. Nature Neuroscience, 18(11), 1664–1671. 10.1038/nn.4135

Fischer-Baum, S., Kook, J. H., Lee, Y., Ramos-Nuñez, A., & Vannucci, M. (2018). Individual Differences in the Neural and Cognitive Mechanisms of Single Word Reading. Front Hum Neurosci, 12, 271. 10.3389/fnhum.2018.00271

Fox, M. D., & Raichle, M. E. (2007). Spontaneous fluctuations in brain activity observed with functional magnetic resonance imaging. Nat Rev Neurosci, 8(9), 700–711. 10.1038/nrn2201

Fox, M. D., Snyder, A. Z., Vincent, J. L., Corbetta, M., Van Essen, D. C., & Raichle, M. E. (2005). The human brain is intrinsically organized into dynamic, anticorrelated functional networks. Proc Natl Acad Sci U S A, 102(27), 9673–9678. 10.1073/pnas.0504136102

Gayet, S., Paffen, C. L. E., & Van der Stigchel, S. (2018). Visual Working Memory Storage Recruits Sensory Processing Areas. Trends in Cognitive Sciences, 22(3), 189–190. 10.1016/j.tics.2017.09.011

Geerligs, L., Rubinov, M., Cam, C., & Henson, R. N. (2015). State and Trait Components of Functional Connectivity: Individual Differences Vary with Mental State. J Neurosci, 35(41), 13949–13961. 10.1523/jneurosci.1324-15.2015

Gelman, A., Goodrich, B., Gabry, J., & Vehtari, A. (2019). R-squared for Bayesian Regression Models. The American Statistician, 73(3), 307–309. 10.1080/00031305.2018.1549100

Genon, S., Eickhoff, S. B., & Kharabian, S. (2022). Linking interindividual variability in brain structure to behaviour. Nature Reviews Neuroscience, 23(5), 307–318. 10.1038/s41583-022-00584-7

Grady, C. (2012). The cognitive neuroscience of ageing. Nature Reviews Neuroscience, 13(7), 491–505. 10.1038/nrn3256

Gratton, C., Laumann, T. O., Nielsen, A. N., Greene, D. J., Gordon, E. M., Gilmore, A. W., Nelson, S. M., Coalson, R. S., Snyder, A. Z., Schlaggar, B. L., Dosenbach, N. U. F., & Petersen, S. E. (2018). Functional Brain Networks Are Dominated by Stable Group and Individual Factors, Not Cognitive or Daily Variation. Neuron, 98(2), 439–452.e435. 10.1016/j.neuron.2018.03.035

Hedden, T., Schultz, A. P., Rieckmann, A., Mormino, E. C., Johnson, K. A., Sperling, R. A., & Buckner, R. L. (2016). Multiple Brain Markers are Linked to Age-Related Variation in Cognition. Cereb Cortex, 26(4), 1388–1400. 10.1093/cercor/bhu238

Herbet, G., Zemmoura, I., & Duffau, H. (2018). Functional Anatomy of the Inferior Longitudinal Fasciculus: From Historical Reports to Current Hypotheses. Front Neuroanat, 12, 77. 10.3389/fnana.2018.00077

Huang, A. S., Klein, D. N., & Leung, H.-C. (2016). Load-related brain activation predicts spatial working memory performance in youth aged 9–12 and is associated with executive function at earlier ages. Developmental Cognitive Neuroscience, 17, 1–9. 10.1016/j.dcn.2015.10.007

Kalpouzos, G., Persson, J., & Nyberg, L. (2012). Local brain atrophy accounts for functional activity differences in normal aging. Neurobiol Aging, 33(3), 623.e621-623.e613. 10.1016/j.neurobiolaging.2011.02.021

Kanai, R., & Rees, G. (2011). The structural basis of inter-individual differences in human behaviour and cognition. Nature Reviews Neuroscience, 12(4), 231–242. 10.1038/nrn3000

Kelso, J. A. (2012). Multistability and metastability: understanding dynamic coordination in the brain. Philos Trans R Soc Lond B Biol Sci, 367(1591), 906–918. 10.1098/rstb.2011.0351

Keogh, R., Wicken, M., & Pearson, J. (2021). Visual working memory in aphantasia: Retained accuracy and capacity with a different strategy. Cortex, 143, 237–253. 10.1016/j.cortex.2021.07.012

Kherif, F., Josse, G., Seghier, M. L., & Price, C. J. (2009). The main sources of intersubject variability in neuronal activation for reading aloud. J Cogn Neurosci, 21(4), 654–668. 10.1162/jocn.2009.21084

King, M., Hernandez-Castillo, C. R., Poldrack, R. A., Ivry, R. B., & Diedrichsen, J. (2019). Functional boundaries in the human cerebellum revealed by a multi-domain task battery. Nature Neuroscience, 22(8), 1371–1378. 10.1038/s41593-019-0436-x

Kraemer, D. J., Schinazi, V. R., Cawkwell, P. B., Tekriwal, A., Epstein, R. A., & Thompson-Schill, S. L. (2017). Verbalizing, visualizing, and navigating: The effect of strategies on encoding a large-scale virtual environment. J Exp Psychol Learn Mem Cogn, 43(4), 611–621. 10.1037/xlm0000314

Lancichinetti, A., & Fortunato, S. (2012). Consensus clustering in complex networks. Scientific Reports, 2(1), 336. 10.1038/srep00336

Lee, L., Siebner, H. R., Rowe, J. B., Rizzo, V., Rothwell, J. C., Frackowiak, R. S., & Friston, K. J. (2003). Acute remapping within the motor system induced by low-frequency repetitive transcranial magnetic stimulation. Journal of Neuroscience, 23(12), 5308–5318.

Lewis-Peacock, J. A., Drysdale, A. T., & Postle, B. R. (2014). Neural Evidence for the Flexible Control of Mental Representations. Cerebral Cortex, 25(10), 3303–3313. 10.1093/cercor/bhu130

Lindenberger, U. (2014). Human cognitive aging: Corriger la fortune? Science, 346(6209), 572–578. doi:10.1126/science.1254403

Linke, A. C., Vicente-Grabovetsky, A., Mitchell, D. J., & Cusack, R. (2011). Encoding strategy accounts for individual differences in change detection measures of VSTM. Neuropsychologia, 49(6), 1476–1486. 10.1016/j.neuropsychologia.2010.11.034

Lövdén, M., Karalija, N., Andersson, M., Wåhlin, A., Axelsson, J., Köhncke, Y., Jonasson, L. S., Rieckman, A., Papenberg, G., Garrett, D. D., Guitart-Masip, M., Salami, A., Riklund, K., Bäckman, L., Nyberg, L., & Lindenberger, U. (2018). Latent-Profile Analysis Reveals Behavioral and Brain Correlates of Dopamine-Cognition Associations. Cereb Cortex, 28(11), 3894–3907. 10.1093/cercor/bhx253

Lucas, M., Wagshul, M. E., Izzetoglu, M., & Holtzer, R. (2018). Moderating Effect of White Matter Integrity on Brain Activation During Dual-Task Walking in Older Adults. The Journals of Gerontology: Series A, 74(4), 435–441. 10.1093/gerona/gly131

Luck, S. J., & Vogel, E. K. (2013). Visual working memory capacity: from psychophysics and neurobiology to individual differences. Trends Cogn Sci, 17(8), 391–400. 10.1016/j.tics.2013.06.006

Lugtmeijer, S., Geerligs, L., Tsvetanov, K. A., Mitchell, D. J., Cam, C. A. N., & Campbell, K. L. (2023). Lifespan differences in visual short-term memory load-modulated functional connectivity. Neuroimage, 270, 119982. 10.1016/j.neuroimage.2023.119982

Madden, D. J., Bennett, I. J., & Song, A. W. (2009). Cerebral White Matter Integrity and Cognitive Aging: Contributions from Diffusion Tensor Imaging. Neuropsychology Review, 19(4), 415–435. 10.1007/s11065-009-9113-2

Magen, H. (2017). The role of central attention in retrieval from visual short-term memory. Psychonomic Bulletin & Review, 24(2), 423–430. 10.3758/s13423-016-1111-9

Mason, P. H., Winter, B., & Grignolio, A. (2015). Hidden in plain view: degeneracy in complex systems. Biosystems, 128, 1–8.

Mill, R. D., & Cole, M. W. (2023). Neural representation dynamics reveal computational principles of cognitive task learning. bioRxiv, 2023.2006.2027.546751. 10.1101/2023.06.27.546751

Mill, R. D., Gordon, B. A., Balota, D. A., & Cole, M. W. (2020). Predicting dysfunctional age-related task activations from resting-state network alterations. NeuroImage, 221, 117167. 10.1016/j.neuroimage.2020.117167

Mill, R. D., Hamilton, J. L., Winfield, E. C., Lalta, N., Chen, R. H., & Cole, M. W. (2022). Network modeling of dynamic brain interactions predicts emergence of neural information that supports human cognitive behavior. PLOS Biology, 20(8), e3001686. 10.1371/journal.pbio.3001686

Miller, M. B., Donovan, C.-L., Bennett, C. M., Aminoff, E. M., & Mayer, R. E. (2012). Individual differences in cognitive style and strategy predict similarities in the patterns of brain activity between individuals. Neuroimage, 59(1), 83–93. 10.1016/j.neuroimage.2011.05.060

Mitchell, D. J., & Cusack, R. (2018). Visual short-term memory through the lifespan: Preserved benefits of context and metacognition. Psychol Aging, 33(5), 841–854. 10.1037/pag0000265

Mitchell, D. J., Mousley, A. L. S., Shafto, M. A., Cam-CAN, & Duncan, J. (2023). Neural Contributions to Reduced Fluid Intelligence across the Adult Lifespan. The Journal of Neuroscience, 43(2), 293–307. 10.1523/jneurosci.0148-22.2022

Morcom, A. M., & Henson, R. N. A. (2018). Increased Prefrontal Activity with Aging Reflects Nonspecific Neural Responses Rather than Compensation. J Neurosci, 38(33), 7303–7313. 10.1523/jneurosci.1701-17.2018

Nagel, I. E., Preuschhof, C., Li, S.-C., Nyberg, L., Bäckman, L., Lindenberger, U., & Heekeren, H. R. (2009). Performance level modulates adult age differences in brain activation during spatial working memory. Proceedings of the National Academy of Sciences, 106(52), 22552–22557. doi:10.1073/pnas.0908238106

Noppeney, U., Friston, K. J., & Price, C. J. (2004). Degenerate neuronal systems sustaining cognitive functions. J Anat, 205(6), 433–442. 10.1111/j.0021-8782.2004.00343.x

Noppeney, U., Penny, W. D., Price, C. J., Flandin, G., & Friston, K. J. (2006). Identification of degenerate neuronal systems based on intersubject variability. Neuroimage, 30(3), 885–890. 10.1016/j.neuroimage.2005.10.010

Noppeney, U., Price, C. J., Penny, W. D., & Friston, K. J. (2005). Two Distinct Neural Mechanisms for Category-selective Responses. Cerebral Cortex, 16(3), 437–445. 10.1093/cercor/bhi123

Nudo, R. (2013). Recovery after brain injury: mechanisms and principles [Review]. Frontiers in Human Neuroscience, 7. 10.3389/fnhum.2013.00887

Nyberg, L., Boraxbekk, C.-J., Sörman, D. E., Hansson, P., Herlitz, A., Kauppi, K., Ljungberg, J. K., Lövheim, H., Lundquist, A., Adolfsson, A. N., Oudin, A., Pudas, S., Rönnlund, M., Stiernstedt, M., Sundström, A., & Adolfsson, R. (2020). Biological and environmental predictors of heterogeneity in neurocognitive ageing: Evidence from Betula and other longitudinal studies. Ageing Research Reviews, 64, 101184. 10.1016/j.arr.2020.101184

Nyberg, L., Lövdén, M., Riklund, K., Lindenberger, U., & Bäckman, L. (2012). Memory aging and brain maintenance. Trends Cogn Sci, 16(5), 292–305. 10.1016/j.tics.2012.04.005

Olson, I. R., Von Der Heide, R. J., Alm, K. H., & Vyas, G. (2015). Development of the uncinate fasciculus: Implications for theory and developmental disorders. Dev Cogn Neurosci, 14, 50–61. 10.1016/j.dcn.2015.06.003

Oschwald, J., Guye, S., Liem, F., Rast, P., Willis, S., Röcke, C., Jäncke, L., Martin, M., & Mérillat, S. (2020). Brain structure and cognitive ability in healthy aging: a review on longitudinal correlated change. Reviews in the Neurosciences, 31(1), 1–57. doi:10.1515/revneuro-2018-0096

Park, D. C., & Reuter-Lorenz, P. (2009). The Adaptive Brain: Aging and Neurocognitive Scaffolding. Annual Review of Psychology, 60(Volume 60, 2009), 173-196. 10.1146/annurev.psych.59.103006.093656

Park, H.-J., & Friston, K. (2013). Structural and Functional Brain Networks: From Connections to Cognition. Science, 342(6158), 1238411. doi:10.1126/science.1238411

Pascual-Leone, A., Amedi, A., Fregni, F., & Merabet, L. B. (2005). The plastic human brain cortex. Annual Review of Neuroscience, 28(1), 377–401. 10.1146/annurev.neuro.27.070203.144216

Patel, R., Mackay, C. E., Jansen, M. G., Devenyi, G. A., O’Donoghue, M. C., Kivimaki, M., Singh-Manoux, A., Zsoldos, E., Ebmeier, K. P., Chakravarty, M. M., & Suri, S. (2022). Inter- and intra-individual variation in brain structural-cognition relationships in aging. Neuroimage, 257, 119254. 10.1016/j.neuroimage.2022.119254

Pearson, J., & Keogh, R. (2019). Redefining Visual Working Memory: A Cognitive-Strategy, Brain-Region Approach. Current Directions in Psychological Science, 28(3), 266–273. 10.1177/0963721419835210

Poldrack, R. A. (2006). Can cognitive processes be inferred from neuroimaging data? Trends in Cognitive Sciences, 10(2), 59–63. 10.1016/j.tics.2005.12.004

Poldrack, R. A., Huckins, G., & Varoquaux, G. (2020). Establishment of Best Practices for Evidence for Prediction: A Review. JAMA Psychiatry, 77(5), 534–540. 10.1001/jamapsychiatry.2019.3671

Postle, B. R. (2015). The cognitive neuroscience of visual short-term memory. Current Opinion in Behavioral Sciences, 1, 40–46. 10.1016/j.cobeha.2014.08.004

Power, J. D., Cohen, A. L., Nelson, S. M., Wig, G. S., Barnes, K. A., Church, J. A., Vogel, A. C., Laumann, T. O., Miezin, F. M., Schlaggar, B. L., & Petersen, S. E. (2011). Functional network organization of the human brain. Neuron, 72(4), 665–678. 10.1016/j.neuron.2011.09.006

Price, C. J., & Friston, K. J. (2002). Degeneracy and cognitive anatomy. Trends in Cognitive Sciences, 6(10), 416–421.

Ritakallio, L., Fellman, D., Salmi, J., Jylkkä, J., & Laine, M. (2024). Self-reported strategy use in working memory tasks. Scientific Reports, 14(1), 4893. 10.1038/s41598-024-54160-3

Rose, N. S. (2020). The Dynamic-Processing Model of Working Memory. Current Directions in Psychological Science, 29(4), 378–387. 10.1177/0963721420922185

Rubinov, M., & Sporns, O. (2010). Complex network measures of brain connectivity: uses and interpretations. Neuroimage, 52(3), 1059–1069. 10.1016/j.neuroimage.2009.10.003

Rubinov, M., & Sporns, O. (2011). Weight-conserving characterization of complex functional brain networks. NeuroImage, 56(4), 2068–2079. 10.1016/j.neuroimage.2011.03.069

Salami, A., Eriksson, J., & Nyberg, L. (2012). Opposing effects of aging on large-scale brain systems for memory encoding and cognitive control. J Neurosci, 32(31), 10749–10757. 10.1523/jneurosci.0278-12.2012

Salami, A., Garrett, D. D., Wåhlin, A., Rieckmann, A., Papenberg, G., Karalija, N., Jonasson, L., Andersson, M., Axelsson, J., Johansson, J., Riklund, K., Lövdén, M., Lindenberger, U., Bäckman, L., & Nyberg, L. (2019).Dopamine D 2/3 Binding Potential Modulates Neural Signatures of Working Memory in a Load-Dependent Fashion. The Journal of Neuroscience, 39(3), 537-547. 10.1523/jneurosci.1493-18.2018

Salami, A., Rieckmann, A., Fischer, H., & Bäckman, L. (2014). A multivariate analysis of age-related differences in functional networks supporting conflict resolution. Neuroimage, 86, 150–163. 10.1016/j.neuroimage.2013.08.002

Salami, A., Rieckmann, A., Karalija, N., Avelar-Pereira, B., Andersson, M., Wåhlin, A., Papenberg, G., Garrett, D. D., Riklund, K., Lövdén, M., Lindenberger, U., Bäckman, L., & Nyberg, L. (2018). Neurocognitive Profiles of Older Adults with Working-Memory Dysfunction. Cerebral Cortex, 28(7), 2525–2539. 10.1093/cercor/bhy062

Samu, D., Campbell, K. L., Tsvetanov, K. A., Shafto, M. A., & Tyler, L. K. (2017). Preserved cognitive functions with age are determined by domain-dependent shifts in network responsivity. Nature communications, 8(1), 14743.

Sander, M. C., Lindenberger, U., & Werkle-Bergner, M. (2012). Lifespan age differences in working memory: A two-component framework. Neuroscience & Biobehavioral Reviews, 36(9), 2007–2033. 10.1016/j.neubiorev.2012.06.004

Sanfratello, L., Caprihan, A., Stephen, J. M., Knoefel, J. E., Adair, J. C., Qualls, C., Lundy, S. L., & Aine, C. J. (2014). Same task, different strategies: How brain networks can be influenced by memory strategy. Human Brain Mapping, 35(10), 5127–5140. 10.1002/hbm.22538

Scimeca, J. M., Kiyonaga, A., & D’Esposito, M. (2018). Reaffirming the Sensory Recruitment Account of Working Memory. Trends in Cognitive Sciences, 22(3), 190–192. 10.1016/j.tics.2017.12.007

Scrucca, L., Fop, M., Murphy, T. B., & Raftery, A. E. (2016). mclust 5: Clustering, Classification and Density Estimation Using Gaussian Finite Mixture Models. R j, 8(1), 289–317.

Seeley, W. W., Menon, V., Schatzberg, A. F., Keller, J., Glover, G. H., Kenna, H., Reiss, A. L., & Greicius, M. D. (2007). Dissociable Intrinsic Connectivity Networks for Salience Processing and Executive Control. The Journal of Neuroscience, 27(9), 2349–2356. 10.1523/jneurosci.5587-06.2007

Seghier, M. L., Lee, H. L., Schofield, T., Ellis, C. L., & Price, C. J. (2008). Inter-subject variability in the use of two different neuronal networks for reading aloud familiar words. Neuroimage, 42(3), 1226–1236. 10.1016/j.neuroimage.2008.05.029

Seghier, M. L., & Price, C. J. (2016). Visualising inter-subject variability in fMRI using threshold-weighted overlap maps. Sci Rep, 6, 20170. 10.1038/srep20170

Seghier, M. L., & Price, C. J. (2018). Interpreting and Utilising Intersubject Variability in Brain Function. Trends in Cognitive Sciences, 22(6), 517–530. 10.1016/j.tics.2018.03.003

Serences, J. T. (2016). Neural mechanisms of information storage in visual short-term memory. Vision Res, 128, 53–67. 10.1016/j.visres.2016.09.010

Shafto, M. A., Tyler, L. K., Dixon, M., Taylor, J. R., Rowe, J. B., Cusack, R., Calder, A. J., Marslen-Wilson, W. D., Duncan, J., Dalgleish, T., Henson, R. N., Brayne, C., & Matthews, F. E. (2014). The Cambridge Centre for Ageing and Neuroscience (Cam-CAN) study protocol: a cross-sectional, lifespan, multidisciplinary examination of healthy cognitive ageing. BMC Neurol, 14, 204. 10.1186/s12883-014-0204-1

Shekari, E., & Nozari, N. (2023). A narrative review of the anatomy and function of the white matter tracts in language production and comprehension. Front Hum Neurosci, 17, 1139292. 10.3389/fnhum.2023.1139292

Smith, S. M., Fox, P. T., Miller, K. L., Glahn, D. C., Fox, P. M., Mackay, C. E., Filippini, N., Watkins, K. E., Toro, R., Laird, A. R., & Beckmann, C. F. (2009). Correspondence of the brain’s functional architecture during activation and rest. Proc Natl Acad Sci U S A, 106(31), 13040–13045. 10.1073/pnas.0905267106

Spurk, D., Hirschi, A., Wang, M., Valero, D., & Kauffeld, S. (2020). Latent profile analysis: A review and “how to” guide of its application within vocational behavior research. Journal of Vocational Behavior, 120, 103445. 10.1016/j.jvb.2020.103445

Stokes, M. G. (2015). &#x2018;Activity-silent&#x2019; working memory in prefrontal cortex: a dynamic coding framework. Trends in Cognitive Sciences, 19(7), 394–405. 10.1016/j.tics.2015.05.004

Tavor, I., Parker Jones, O., Mars, R. B., Smith, S. M., Behrens, T. E., & Jbabdi, S. (2016). Task-free MRI predicts individual differences in brain activity during task performance. Science, 352(6282), 216–220. 10.1126/science.aad8127

Taylor, J. R., Williams, N., Cusack, R., Auer, T., Shafto, M. A., Dixon, M., Tyler, L. K., Cam, C., & Henson, R. N. (2017). The Cambridge Centre for Ageing and Neuroscience (Cam-CAN) data repository: Structural and functional MRI, MEG, and cognitive data from a cross-sectional adult lifespan sample. Neuroimage, 144(Pt B), 262–269. 10.1016/j.neuroimage.2015.09.018

Traag, V. A., Waltman, L., & van Eck, N. J. (2019). From Louvain to Leiden: guaranteeing well-connected communities. Scientific Reports, 9(1), 5233. 10.1038/s41598-019-41695-z

Van Essen, D. C., Drury, H. A., Dickson, J., Harwell, J., Hanlon, D., & Anderson, C. H. (2001). An integrated software suite for surface-based analyses of cerebral cortex. Journal of the American Medical Informatics Association, 8(5), 443–459.

Vermunt, J. K., & Magidson, J. (2002). Latent class cluster analysis. Applied latent class analysis, 11(89-106), 60.

Von Der Heide, R. J., Skipper, L. M., Klobusicky, E., & Olson, I. R. (2013). Dissecting the uncinate fasciculus: disorders, controversies and a hypothesis. Brain, 136(Pt 6), 1692–1707. 10.1093/brain/awt094

Walhovd, K. B., Lövden, M., & Fjell, A. M. (2023). Timing of lifespan influences on brain and cognition. Trends in Cognitive Sciences, 27(10), 901–915. 10.1016/j.tics.2023.07.001

Westlin, C., Theriault, J. E., Katsumi, Y., Nieto-Castanon, A., Kucyi, A., Ruf, S. F., Brown, S. M., Pavel, M., Erdogmus, D., & Brooks, D. H. (2023). Improving the study of brain-behavior relationships by revisiting basic assumptions. Trends in Cognitive Sciences.

Wig, G. S., Schlaggar, B. L., & Petersen, S. E. (2011). Concepts and principles in the analysis of brain networks. Annals of the New York Academy of Sciences, 1224(1), 126–146. 10.1111/j.1749-6632.2010.05947.x

Yeo, B. T., Krienen, F. M., Sepulcre, J., Sabuncu, M. R., Lashkari, D., Hollinshead, M., Roffman, J. L., Smoller, J. W., Zöllei, L., Polimeni, J. R., Fischl, B., Liu, H., & Buckner, R. L. (2011). The organization of the human cerebral cortex estimated by intrinsic functional connectivity. J Neurophysiol, 106(3), 1125–1165. 10.1152/jn.00338.2011

Zeman, A. (2024). Aphantasia and hyperphantasia: exploring imagery vividness extremes. Trends in Cognitive Sciences, 28(5), 467–480. 10.1016/j.tics.2024.02.007

Zuo, N., Salami, A., Yang, Y., Yang, Z., Sui, J., & Jiang, T. (2019). Activation-based association profiles differentiate network roles across cognitive loads. Human Brain Mapping, 40(9), 2800–2812. 10.1002/hbm.24561

