## Supplemental for "Multiple brain activation patterns underlying successful visual short-term memory across the adult lifespan"

### Supplementary figures

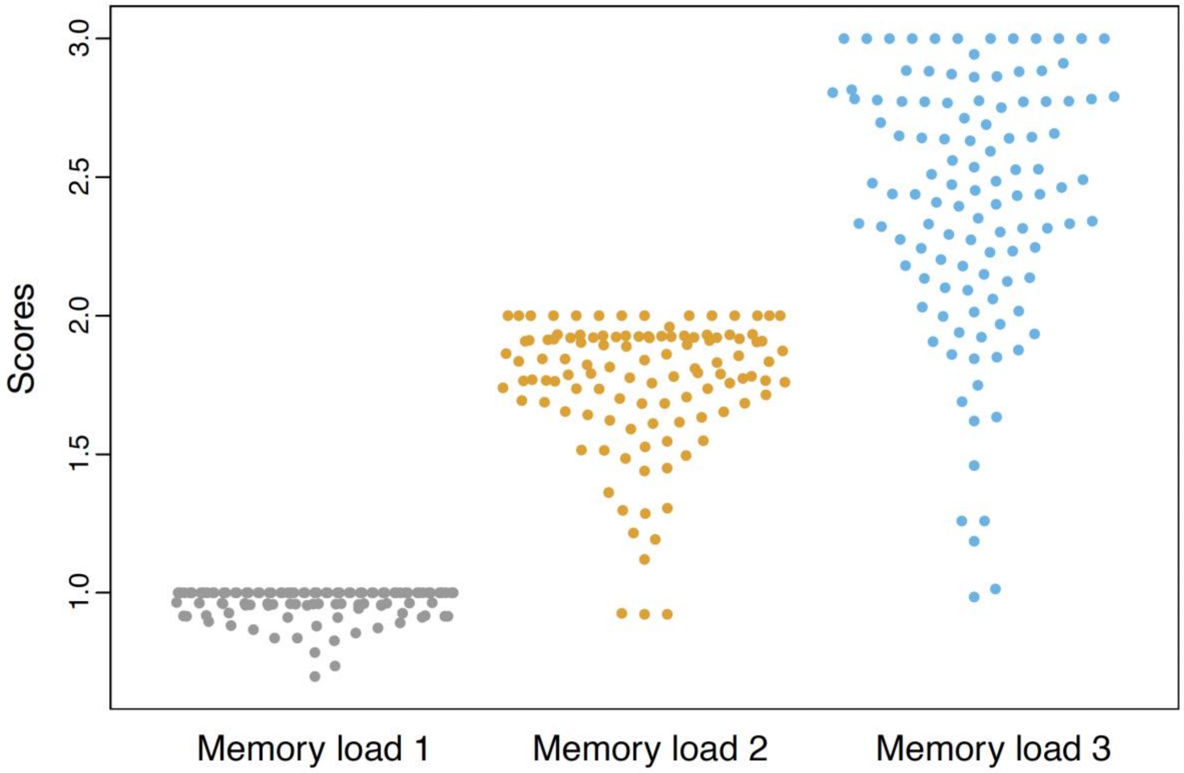

**Supplementary Figure 1:** The distribution of the number of items in memory at each memory load (load 1-3) as illustrated with beeswarm plots. The number of items in memory for load 1 displayed a ceiling effect, where many participants achieved high performance. This was also present to some extent at load 2. Therefore, the analyses in this manuscript focused on load 3, where the data demonstrated more variation.

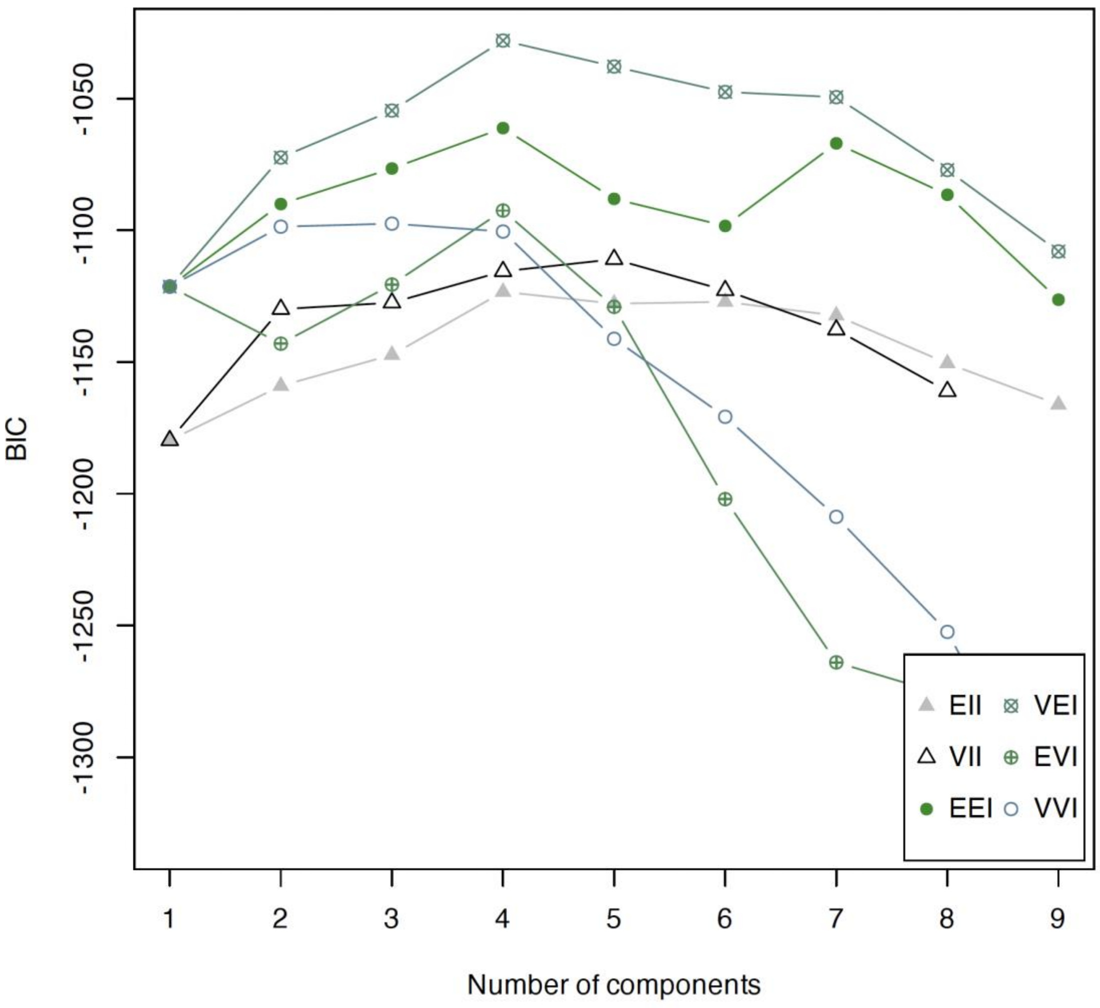

**Supplementary Figure 2**: Distribution of Bayesian Information Criterion (BIC) values for the different latent-profile models (1-5 clusters) estimated based on the residual activity in each brain network as well as the overall responsivity across all networks. The most optimal BIC value (BIC = -1028) corresponded to a model with 4 latent subgroups of equal shape, variable volume, and a diagonal distribution ("VEI").

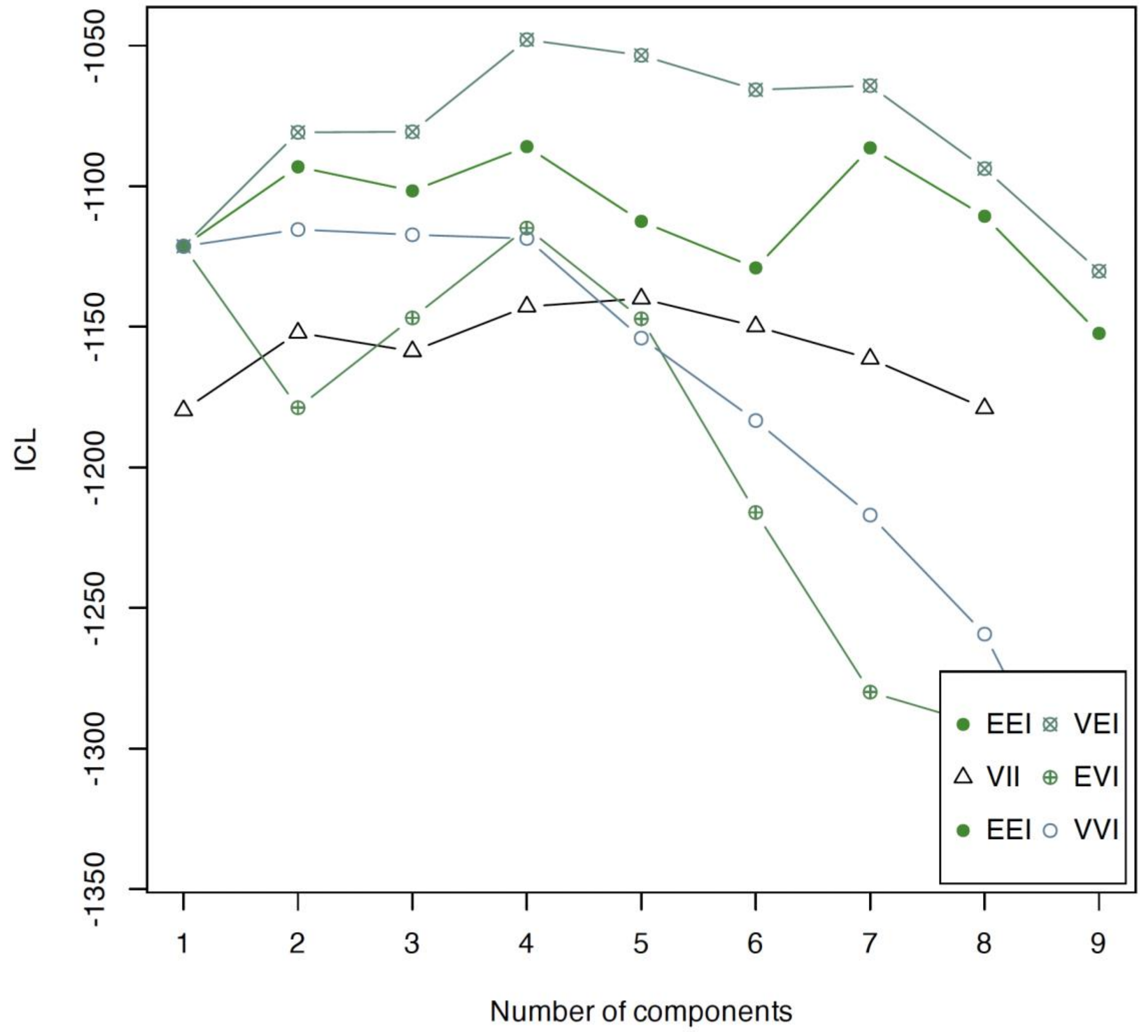

**Supplementary Figure 3**: Distribution of Integrated Completed Likelihood (ICL) values for the different latent-profile models (1-5 clusters) estimated based on the residual activity in each brain network as well as the overall responsivity across all networks. The most optimal ICL value (ICL = -1048) corresponded to a model with 4 latent subgroups of equal shape, variable volume, and a diagonal distribution (“VEI”).

### Supplementary tables

**Supplementary Table 1. Model fit indices latent profile analysis**

| Number of classes | df | BIC | ICL | LRT | p-value |
|---|---|---|---|---|---|
| 1 | 16 | -1121.36 | -1121.36 | | |
| 2 | 26 | -1072.37 | -1080.84 | 96.26 | 0.001 |
| 3 | 36 | -1054.57 | -1080.66 | 65.08 | 0.001 |
| 4 | 46 | **-1028.00** | **-1047.89** | 73.85 | 0.001 |
| 5 | 56 | -1037.85 | -1054.45 | 37.42 | 0.003 |
| 6 | 66 | -1047.51 | -1065.71 | 37.61 | 0.009 |
| 7 | 76 | -1049.46 | -1064.24 | 45.33 | 0.001 |
| 8 | 86 | -1077.12 | -1093.73 | 19.61 | **0.422** |

BIC = Bayesian Information Criterion; ICL = Integrated Completed Likelihood; LRT = Likelihood Ratio Test.

**Supplementary Table 2. Between-group comparisons of brain module activity**

| | | No correction applied | | | | | Corrected for age | | | | |
|---|---|---|---|---|---|---|---|---|---|---|---|
| Brain module | N | F-statistic | p-value | q-value | BF | $R^2$ | F-statistic | p-value | q-value | BF | $R^2$ |
| **FCM** | **113** | **54.29*** | **< 0.001** | **< 0.001** | **3.00e+14** | **0.51†** | **36.92*** | **< 0.001** | **< 0.001** | **3.52e+12** | **0.46†** |
| TPM | 113 | 1.05 | 0.383 | 0.44 | 1.00e-01 | 0.00 | 36.79 | 0.425 | 0.4859 | 1.27e-01 | 0.00 |
| CTM | 113 | 10.92* | < 0.001 | < 0.001 | 1.59e+02 | 0.15 | 35.76* | < 0.001 | < 0.001 | 1.78e+02 | 0.15 |
| **VM** | **113** | **33.93*** | **< 0.001** | **< 0.001** | **2.80e+12** | **0.46†** | **37.23*** | **< 0.001** | **< 0.001** | **4.26e+11** | **0.44†** |
| **DAM** | **113** | **20.00*** | **< 0.001** | **< 0.001** | **8.16e+05** | **0.28†** | **36.58*** | **< 0.001** | **< 0.001** | **6.37e+04** | **0.24** |
| MM | 113 | 9.45* | < 0.001 | < 0.001 | 2.36e+02 | 0.15 | 36.38* | < 0.001 | <0.001 | 1.27e+02 | 0.14 |
| **DMM** | **113** | **42.56*** | **< 0.001** | **< 0.001** | **1.38e+13** | **0.47†** | **37.13*** | **< 0.001** | **< 0.001** | **4.72e+11** | **0.44†** |
| Resp | 113 | 0.77 | 0.517 | 0.517 | 7.00e-02 | 0.00 | 38.19 | 0.512 | 0.512 | 6.40e-02 | 0.00 |

Results from the Welch's ANOVA and corresponding Bayes Factor (BF) for the alternative hypothesis and Bayesian $R^2$. Results were considered as significant if both FDR-corrected q-value < 0.05 and BF > 3. Differences were most pronounced for the frontal control module (FCM), visual module (VM), dorsal attention module (DAM), and default mode module (DMM), as reflected by the corresponding values of BF and Bayesian $R^2$. Results were not affected by age corrections. BF = Bayes Factor; FCM = frontal control module; TPM = temporal parietal module; CTM = cerebellum/thalamus module; VM = visual module; DAM = dorsal attention module; MM = motor module; DMM = default mode module.

*= FDR-corrected q-value < 0.001 & BF > 100.

†$R^2 \geq 0.26$.

**Supplementary Table 3. Between-group comparisons of participant characteristics**

| Continuous variables | N | Df | Df error | F-statistic | p-value | BF | R2 |
|---|---|---|---|---|---|---|---|
| Age | 113 | 3 | 40.77 | 2.79 | 0.05 | 0.43 | 0.00 |
| Number of items in memory (load 3) | 113 | 3 | 44.12 | 0.45 | 0.72 | 0.04 | 0.00 |
| Memory precision (load 3) | 113 | 3 | 43.36 | 1.98 | 0.13 | 0.18 | 0.00 |
| Categorical variables | N | Df | $\chi^2$-statistic | 95% CI | p-value | BF | Cramer's V |
| Sex | 113 | 3 | 3.43 | 0.00-1.00 | 0.32 | 0.16 | 0.07 |
| Educational attainment | 113 | 3 | 4.28 | 0.00-0.30 | 0.23 | 0.26 | 0.11 |

Upper part displays the results from the Welch's ANOVA and corresponding Bayes Factor (BF) for the alternative hypothesis and Bayesian $R^2$. Lower part displays results from the Chi-squared test and corresponding Bayes Factor (BF) for the alternative hypothesis. Results were considered as significant if both FDR-corrected q-value < 0.05 and BF > 3. Notably, educational attainment was statistically compared after dichotomizing the four corresponding categories (university as highest level education vs. the other catgories; A' levels, GCSE grade, none above 16). The between-group comparisons of memory performance were not affected by age corrections.

BF = Bayes Factor.

**Supplementary Table 4. Within-group associations between task performance and brain module activity**

| | Whole sample | | | | Subgroup 1 | | | | Subgroup 2 | | | | Subgroup 3 | | | |
|---|---|---|---|---|---|---|---|---|---|---|---|---|---|---|---|---|
| Memory precision (load 3) | *N* = 101 | | | | *N* = 35 | | | | *N* = 24 | | | | *N* = 42 | | | |
| | R [95% CI] | p-value | q-value | BF | R [95% CI] | p-value | q-value | BF | R [95% CI] | p-value | q-value | BF | r [95% CI] | p-value | q-value | BF |
| FCM | -0.15 [-0.33; 0.05] | 0.136 | 0.271 | 0.45 | -0.37 [-0.62; -0.04] | **0.030 *** | 0.059 | 2.36 | -0.41 [-0.70; 0.01] | **0.045 *** | 0.180 | 1.93 | 0.12 [-0.19; 0.41] | 0.465 | 0.630 | 0.30 |
| VM | -0.04 [-0.23; 0.16] | 0.718 | 0.718 | 0.16 | -0.06 [-0.38; 0.28] | 0.742 | 0.742 | 0.27 | 0.28 [-0.14; 0.61] | 0.191 | 0.255 | 0.67 | -0.15 [-0.43; 0.16] | 0.341 | 0.630 | 0.36 |
| DAM | 0.17 [-0.03; 0.35] | 0.098 | 0.271 | 0.58 | 0.42 [0.10; 0.66] | **0.012 *** | **0.048 *** | 4.97 | 0.30 [-0.12; 0.63] | 0.152 | 0.255 | 0.79 | -0.06 [-0.36; 0.25] | 0.696 | 0.696 | 0.25 |
| DMM | 0.06 [-0.14; 0.25] | 0.567 | 0.718 | 0.18 | 0.08 [-0.26; 0.40] | 0.637 | 0.742 | 0.24 | -0.14 [-0.52;0.27] | 0.498 | 0.498 | 0.38 | 0.11 [-0.20; 0.40] | 0.472 | 0.620 | 0.30 |
| Number of items in memory (load 3) | | | | | | | | | | | | | | | | |
| FCM | 0.10 [-0.10; 0.29] | 0.316 | 0.422 | 0.25 | 0.24 [-0.10; 0.53] | 0.167 | 0.333 | 0.63 | 0.17 [-0.24; 0.54] | 0.418 | 0.679 | 0.41 | -0.10 [-0.39; 0.21 | 0.531 | 0.942 | 0.28 |
| VM | 0.03 [-0.17; 0.22] | 0.793 | 0.793 | 0.16 | 0.11 [-0.23; 0.42] | 0.541 | 0.541 | 0.30 | -0.09 [-0.32; 0.42] | 0.679 | 0.679 | 0.33 | -0.02 [-0.32; 0.29] | 0.917 | 0.942 | 0.24 |
| DAM | 0.12 [-0.08; 0.31] | 0.231 | 0.422 | 0.31 | 0.19 [-0.15; 0.50] | 0.264 | 0.352 | 0.46 | 0.18 [-0.24; 0.55] | 0.390 | 0.679 | 0.43 | 0.01 [-0.29; 0.31] | 0.941 | 0.942 | 0.23 |
| DMM | -0.18 [-0.36’ 0.02] | 0.073 | 0.290 | 0.74 | -0.38 [-0.63 – -0.05] | **0.026 *** | 0.102 | 2.67 | -0.11 [-0.50; 0.30 | 0.597 | 0.679 | 0.35 | 0.01 [-0.29; 0.31] | 0.942 | 0.942 | 0.23 |

Pearson correlations the associations between visual short-term memory task performance and brain module activity, for each subgroup and corrected for age (i.e., partial correlations). Confidence intervals are standardized. CI = confidence interval; FCM = frontal control module; VM = visual module; DAM = dorsal attention module; DMM = default mode module.

*= FDR-corrected q-value < 0.05 & BF > 3.

**Supplementary Table 5. Between-group comparisons of grey matter volumes in each brain module**

| | | Corrected for total intracranial volume | | | | | Corrected for total intracranial volume and age | | | | |
|---|---|---|---|---|---|---|---|---|---|---|---|
| Grey matter volume | N | F | p | q | BF | $R^2$ | F | p | q | BF | $R^2$ |
| FCM | 113 | 0.90 | 0.45 | 0.45 | 0.06 | 0.00 | 0.15 | 0.93 | 0.93 | 0.03 | 0.00 |
| TPM | 113 | 5.67** | 0.002 | 0.02 | 5.10 | 0.08 | 2.87* | 0.05 | 0.34 | 0.72 | 0.00 |
| CTM | 113 | 3.54* | 0.02 | 0.08 | 0.89 | 0.00 | 2.04 | 0.12 | 0.43 | 0.11 | 0.00 |
| VM | 113 | 2.97* | 0.04 | 0.08 | 0.57 | 0.00 | 1.04 | 0.39 | 0.67 | 0.07 | 0.00 |
| DAM | 113 | 2.62 | 0.06 | 0.09 | 0.20 | 0.00 | 0.71 | 0.55 | 0.77 | 0.06 | 0.00 |
| MM | 113 | 3.08 | 0.04 | 0.08 | 0.26 | 0.00 | 1.14 | 0.34 | 0.67 | 0.06 | 0.00 |
| DMM | 113 | 1.78 | 0.17 | 0.19 | 0.13 | 0.00 | 0.32 | 0.81 | 0.93 | 0.04 | 0.00 |

Results from the Welch's ANOVA and corresponding Bayes Factor (BF) for the alternative hypothesis and Bayesian $R^2$, corrected for total intracranial volume or additionally for age. To correct for these effects, we regressed the covariate(s) on the variable of interest and used the residuals of that regression. Results were considered as significant if both FDR-corrected q-value < 0.05 and BF > 3. The between-group differences of grey matter volume in the temporal parietal module (TPM) did not remain significant after applying additional age corrections.

BF = Bayes Factor; FCM = frontal control module; TPM = temporal parietal module; CTM = cerebellum/thalamus module; VM = visual module; DAM = dorsal attention module; MM = motor module; DMM = default mode module.

*=uncorrected p-value < 0.05

**=FDR-corrected q-value < 0.05 & BF > 3.

**Supplementary Table 6. Between-group comparisons of mean kurtosis in each white matter tract**

| | | No covariates | | | | | Corrected for age | | | | |
|---|---|---|---|---|---|---|---|---|---|---|---|
| Mean kurtosis | N | F | p-value | q-value | BF | $R^2$ | F | p-value | q-value | BF | $R^2$ |
| Left anterior thalamic radiation | 96 | 2.95 | 0.05 | 0.07 | 0.88 | 0.00 | 1.89 | 0.15 | 0.22 | 0.27 | 0.00 |
| Right anterior thalamic radiation | 96 | 1.80 | 0.16 | 0.21 | 0.22 | 0.00 | 1.28 | 0.30 | 0.41 | 0.10 | 0.00 |
| Left dorsal cingulate gyrus | 96 | 3.57 | 0.02 | 0.06 | 2.69 | 0.07 | 2.69 | 0.06 | 0.22 | 0.92 | 0.00 |
| Right dorsal cingulate gyrus | 95 | 0.71 | 0.55 | 0.58 | 0.07 | 0.00 | 0.49 | 0.69 | 0.73 | 0.05 | 0.00 |
| Left ventral cingulate gyrus | 86 | 0.12 | 0.95 | 0.95 | 0.04 | 0.00 | 0.33 | 0.81 | 0.81 | 0.05 | 0.00 |
| Right ventral cingulate gyrus | 94 | 1.13 | 0.35 | 0.40 | 0.12 | 0.00 | 1.21 | 0.32 | 0.41 | 0.12 | 0.00 |
| Left cerebrospinal tract | 95 | 5.57 ** | 0.00 | 0.02 | 0.84 | 0.00 | 2.06 | 0.12 | 0.22 | 0.15 | 0.00 |
| Right cerebrospinal tract | 94 | 3.18 | 0.04 | 0.07 | 1.37 | 0.04 | 2.81 | 0.05 | 0.22 | 0.56 | 0.00 |
| Forceps major | 96 | 2.73 | 0.06 | 0.09 | 0.80 | 0.00 | 2.13 | 0.11 | 0.22 | 0.32 | 0.00 |
| Forceps minor | 96 | 3.81 | 0.02 | 0.05 | 1.02 | 0.01 | 2.05 | 0.12 | 0.22 | 0.30 | 0.00 |
| Left inferior frontal-occipital fasciculus | 96 | 4.73 *** | 0.01 | 0.03 | 5.04 | 0.09 | 3.02* | 0.04 | 0.22 | 1.16 | 0.02 |
| Right inferior frontal-occipital fasciculus | 96 | 4.08 *** | 0.01 | 0.05 | 3.49 | 0.08 | 2.36 | 0.09 | 0.22 | 0.60 | 0.00 |
| **Left inferior longitudinal fasciculus** | **96** | **6.71 *** ** | **0.00** | **0.00*** | **28.10** | **0.14** | **5.42 *** ** | **0.00** | **0.03** | **5.81** | **0.10** |
| Right inferior longitudinal fasciculus | 95 | 3.40 * | 0.03* | 0.06 | 2.18 | 0.06 | 2.54 | 0.07 | 0.22 | 0.69 | 0.00 |
| Left superior longitudinal fasciculus | 95 | 1.91 | 0.15 | 0.20 | 0.23 | 0.00 | 1.12 | 0.36 | 0.41 | 0.11 | 0.00 |
| Right superior longitudinal fasciculus | 96 | 3.18 | 0.03 | 0.07 | 1.00 | 0.00 | 1.09 | 0.37 | 0.41 | 0.13 | 0.00 |
| **Left uncinate fasciculus** | **96** | **6.60 *** ** | **0.00** | **0.01** | **16.93** | **0.13** | **6.64 *** ** | **0.00** | **0.02** | **17.32** | **0.13** |
| Right uncinate fasciculus | 96 | 1.60 | 0.21 | 0.25 | 0.20 | 0.00 | 1.89 | 0.15 | 0.22 | 0.28 | 0.00 |

Results from the Welch's ANOVA and corresponding Bayes Factor (BF) for the alternative hypothesis and Bayesian $R^2$, with and without additional corrections for age and average mean kurtosis across all tracts. To correct for these effects, we regressed the covariate(s) on the variable of interest and used the residuals of that regression. Results were considered as significant if both FDR-corrected q-value < 0.05 and BF > 3. Only the left inferior longitudinal fasciculus and left uncinate fasciculus demonstrated significant differences between-groups after age corrections. BF = Bayes Factor.

*= uncorrected p-value < 0.05; **=FDR-corrected q-value < 0.05

***=FDR-corrected q-value < 0.05 & BF > 3.
